# An X-linked sex determination mechanism in cannabis and hop

**DOI:** 10.1101/2024.12.09.627636

**Authors:** Sarah B. Carey, Philip C. Bentz, John T. Lovell, Laramie M. Akozbek, Zack A. Myers, Walid Korani, Joshua S. Havill, Lillian Padgitt-Cobb, Ryan C. Lynch, Nicholas Allsing, Jack Mangels, Zachary Stansell, George Stack, Tyler Gordon, Austin Osmanski, Katherine A. Easterling, Leonardo R. Orozco, Zach E. Marcus, Haley Hale, Hannah McCoy, Zachary Meharg, Jane Grimwood, Lawrence B. Smart, Daniela Vergara, Rafael F. Guerrero, Nolan C. Kane, Rich Fletcher, John K. McKay, Todd P. Michael, Gary J. Muehlbauer, Josh Clevenger, Alex Harkess

**Author notes:** Author for correspondence: A.H.

## Abstract

Sex chromosomes in cannabis and hop were identified a century ago because of their obvious visible differences in size (heteromorphy). However, we know little about the genes they contain that control the development of the inflorescences. Here we assembled genomes, with phased sex chromosomes, for hop and cannabis. The XY chromosomes share an origin prior to the divergence between the genera >36 MYA. Due to the inheritance patterns of the XYs, the male-specific region of the Y is highly-degenerated, with substantial gene loss, while the X shows faster rates of molecular evolution. Consistent with the theory that these species lack an active-Y system, no clear sex-determining genes reside on the Y. Instead, an X-linked homolog of aminocyclopropane-1-carboxylate synthase (*ACS*), that is involved in the ethylene biosynthesis pathway, determines the fate of the female inflorescence. Beyond sex determination, the sex chromosomes contribute to the sexual dimorphism in ecology and physiology and have played a role in the domestication and breeding of these species.

## Main Text

The inflorescences in the Cannabaceae family have a deep history of human application. *Cannabis sativa* L. was cultivated by humans around twelve thousand years ago (*1*) and is still grown globally for medicinal compounds (cannabinoids) concentrated in glandular trichomes on female inflorescences (*2*), as well as for multi-purpose oil in seeds, and bast fiber in stems. The female inflorescences of hop (*Humulus lupulus* L.) also concentrate secondary compounds in glandular trichomes and have been used for brewing beer (*3*) since as far back as 1,200 years ago, where they provide the characteristic bitterness and aroma. Thus females in both genera have been selected upon over centuries of human domestication. While there is a chromosomal basis that determines sex in these species, a major complication is that dioecy is not strict. XX females have been shown to produce male flowers, which can cause unintended fertilization and changes in secondary metabolite production (*4*, *5*). Curiously, reversions to monoecy have also been observed in XX and XY individuals, particularly in cannabis (*6*, *7*). Identifying the genes involved with sex-determination is critical to better control leaky sex expression and enhance the production of fiber, medicinal compounds, and oils.

Since their discovery (*8*), sex chromosomes have been of keen interest to study because of their unique inheritance patterns that drive differences in their size, structure, and gene content, as well as their role in sex determination. A century ago, the sex chromosomes of cannabis and hop were among the first to be identified in flowering plants because of their heteromorphic XY pairs (*9*, *10*). *Humulus lupulus* var. *lupulus*, the European hop, has one of the only known XY pairs in plants where the Y chromosome is cytologically smaller than the X. *Cannabis*, instead, has a Y chromosome that is larger than the X, but this form of heteromorphy is also rare across angiosperms; most examined plant sex chromosomes are homomorphic (*11*), including the other botanical varieties of *H. lupulus* (*12*). Although there is extreme variation in cytotype, the shared origin of the XY pairs (*13*) suggests the genera share the same sex determination mechanism.

Here we assemble fully-phased, chromosome-scale genomes with both X and Y chromosomes for cannabis and hop and use these genomes to examine their sex chromosome structure and evolution. Counter to the expectations from other XY systems like in mammals, where the non-recombining region of the Y contains the sex-determining gene (*14*), we instead identify a candidate sex-determination gene on the X, consistent with the leading theory that hop and cannabis do not have an active-Y system (*15*). This result, along with the additional floral genes identified, underline the importance of the sex chromosomes in the development of the economically important cannabis and hop flowers.

### Phased sex chromosome assemblies

Until recently, assembling Cannabaceae genomes was a challenge, because species in the family are highly heterozygous with repeat-rich genomes (*16*, *17*). Further, sex chromosomes are particularly arduous to contiguously assemble (*18*). For example, the Y chromosome was the last to be included in the human ‘T2T’ genome (*19*), and sex chromosomes are incomplete in many mammalian genomes. While progress has been made to identify sex-linked sequences (*20*, *21*), to our knowledge no approach has been designed to assemble and validate perfectly-phased sex chromosomes across the immense range of sizes and levels of divergence that exist (*22*). Therefore, we developed a custom pipeline to phase the sex chromosomes that relies on the generation of male-specific *k*-mers (henceforth, Y-mers (*23*)) and a haplotype-combined scaffolding approach to identify X-versus Y-linked contigs (Supplementary Text).

To examine the structure and gene content of the Cannabaceae sex chromosomes, we built genomes that span the range of XY cytotypes. This includes males of three botanical varieties of hop used across the breeding pedigree (*24*): a line of *H. lupulus* var. *lupulus* (USDA 21110M) and two wild, American relatives, *H. lupulus* var. *lupuloides* and var. *neomexicanus*. We also built genomes for two male cultivars of cannabis (‘Carmagnola’ and ‘Otto II’) and two cultivars that have reverted from dioecy to monoecy (‘USO 31’ and ‘Futura 75’). We used hifiasm with PacBio HiFi long reads and Omni-C incorporation to build contig assemblies (*25*) (Fig. S1-2; Table S1). Prior to scaffolding, we identified X- and Y-linked contigs and manually placed them into separate haplotypes (Table S2; Supplementary Text). After necessary sex chromosome phase correction, in cannabis the haplotype 1 (HAP1) and 2 (HAP2) assemblies are 760.6-828.8 Mb, while the hop genomes are three times larger at 2.4-2.7 Gb (initial contig N50s: 22-167.6 Mb; Table S3). These genomes contain very few gaps (Fig. S3-4) and exhibit Merqury *k*-mer completeness values (*26*) of 93-98.3% and QV values of 38.7-58 (Table S3). Importantly, the sex chromosomes of these assemblies meet the expected cytological patterns. The Y is the largest chromosome in the cannabis genome at 110.6-114 Mb, while the X is 76% smaller at 84.4-87.4 Mb (Fig. 1). Similar to the increased overall genome size, the X chromosomes of the three varieties of hop are three times larger than in cannabis, at 267.4-272 Mb. The hop Y chromosomes show the most drastic difference overall. The American hop Y chromosomes are cytologically homomorphic, where they are 89.3-92% the size of the X. The Y in *H. lupulus* var. *lupulus* is the smallest chromosome in its genome, and the smallest hop Y, at 168.7 Mb fitting the approximate 1.5:1 ratio observed in the cytotype (62.9% the size of the X) (*12*). These near-perfect, phased sex chromosomes mark a methodological advancement in the field of comparative genomics, enabling the generation of chromosome-scale assemblies from a single heterogametic sex individual without parental trio-binning.

**Fig. 1.**
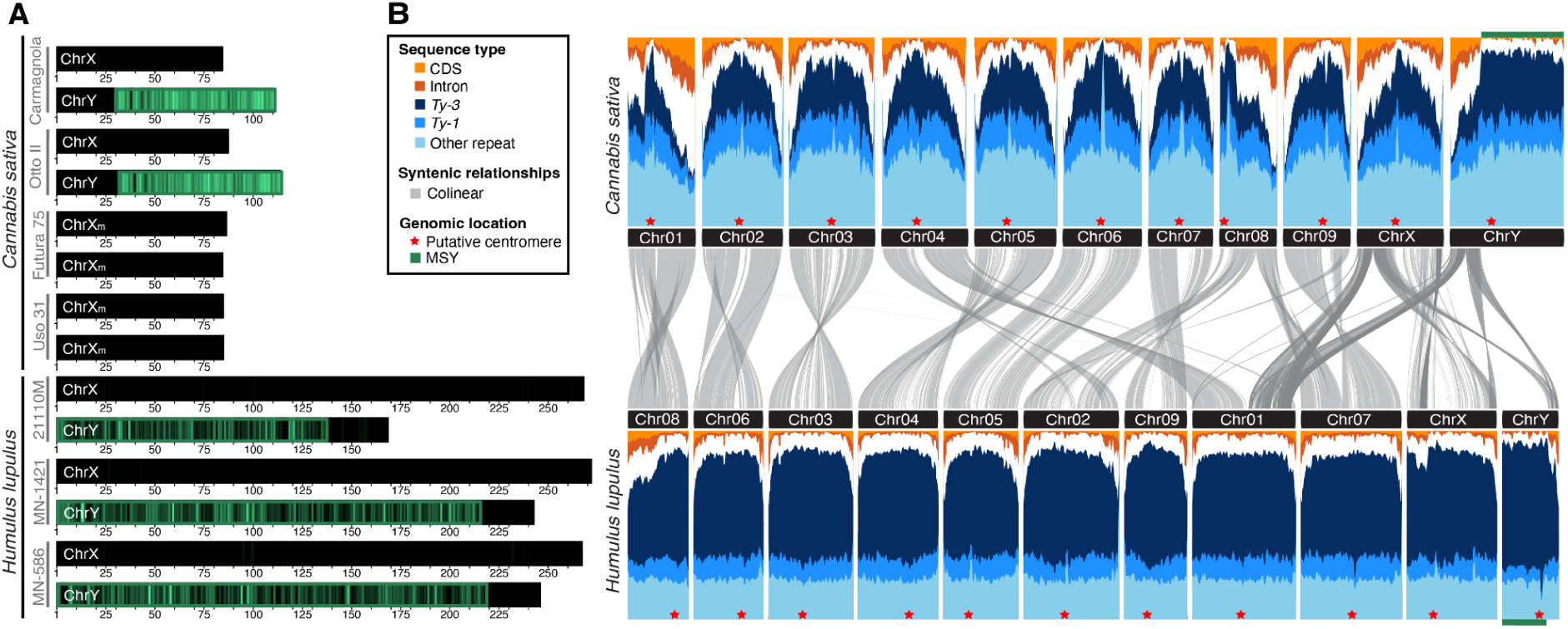
Genome architecture of *Cannabis* and *Humulus*. **A)** Identification of the male-specific region of the Y chromosomes (MSY). The heatmap within each ideogram shows coverage of male-specific *k*-mers within 1Mb windows and a 100kb jump. **B**) gene and repeat landscapes, and syntenic relationships, for cannabis and hop. The chromosomes are represented with a single haplotype for the autosomes but both sex chromosomes. Due to the genome size differences, *Humulus* was scaled to *Cannabis*.

### The origin of cannabis and hop sex chromosomes

In an XY system, the pseudoautosomal region (PAR) recombines relatively freely (*27*), while the male-specific region of the Y (MSY) is typically non-recombining and only inherited by males. Understanding the timing of gene capture into the MSY is important, as the sex-determining genes are likely those that ceased recombining first. Reconstructions of sexual systems suggest that dioecy is the ancestral state in Cannabaceae and its sister families Moraceae and Urticaceae, that contain figs, mulberries, and nettles (*28*). Example species within these families have been shown to switch sex when ethylene concentration or perception is modified (*29*, *30*) , suggesting the possibility of a shared sex-determining mechanism. Currently it is unknown if the two families share an ancestral XY pair; however, recent assemblies in Moraceae (*31*, *32*) and our phased XY chromosomes provide an opportunity to refine our understanding of the timing of the evolution of the sex chromosomes and uncover which genes have the oldest origin.

We used a rigorous Y-mer mapping and gene tree approach to delineate the boundary between the MSY and the PAR. The hop and cannabis Y chromosomes contain a single PAR with a majority of the sequence contained within the MSY (72.4-88.7%), also matching cytological observations (Fig. 1) (*33*). We used a combination of RNAseq and protein homology, and annotated on average 34.9k gene models per haplotype using BRAKER (*34*, *35*), with BUSCOs ranging between 93-97.2% (Table S3-4). Most genes on the Cannabaceae sex chromosomes reside in the PARs and maintain a linear order between XY pairs within and across species, in utter contrast to the MSYs that have very few genes and little synteny remaining (Fig. 1).

To investigate the origin of sex chromosome evolution we used evidence from a suite of complementary analyses. Syntenic relationships show the sex chromosomes are located on the same chromosome in cannabis and hop, but different chromosomes in mulberries and figs (Fig. 2). Likewise, using gene trees we found support for a shared origin between cannabis and hop, where sex-linkage of a gene prior to the divergence of the genera shows all Y-linked copies forming a clade that is sister to the X-linked clade (Fig. S5-S6). No gene tree topology suggests that Moraceace shares this origin. To estimate the age of the MSY, we used synonymous protein changes (Ks) between one-to-one XY orthologs (hereafter, gametologs). Using Ks in hop (max = 0.4; mean top 10% = 0.33), we estimate the sex chromosomes evolved 38.6-48.3 MYA. We next used a phylogeny based on whole-plastome sequences and fossil records to calibrate and estimate a divergence time tree to compare this date to age estimates across Rosales. Resulting age estimates were transposed onto a species tree, inferred using a multi-species coalescent summary approach (*36*) with 775 nuclear gene trees at concordant nodes (Fig. 2, S6). The ancestors of *Cannabis* and *Humulus* began diverging between 33.9-41.4 MYA (95% CI; mean=36.38), whereas Cannabaceae diverged from Moraceae around 66-84.4 MYA (95% CI; mean=74.2). These results firmly place the sex chromosome origin on the stem branch leading to the split between *Cannabis* and *Humulus*, but nonetheless support that their shared XY evolution is among the oldest identified in plants, and independent from the species in Moraceae.

**Fig. 2.**
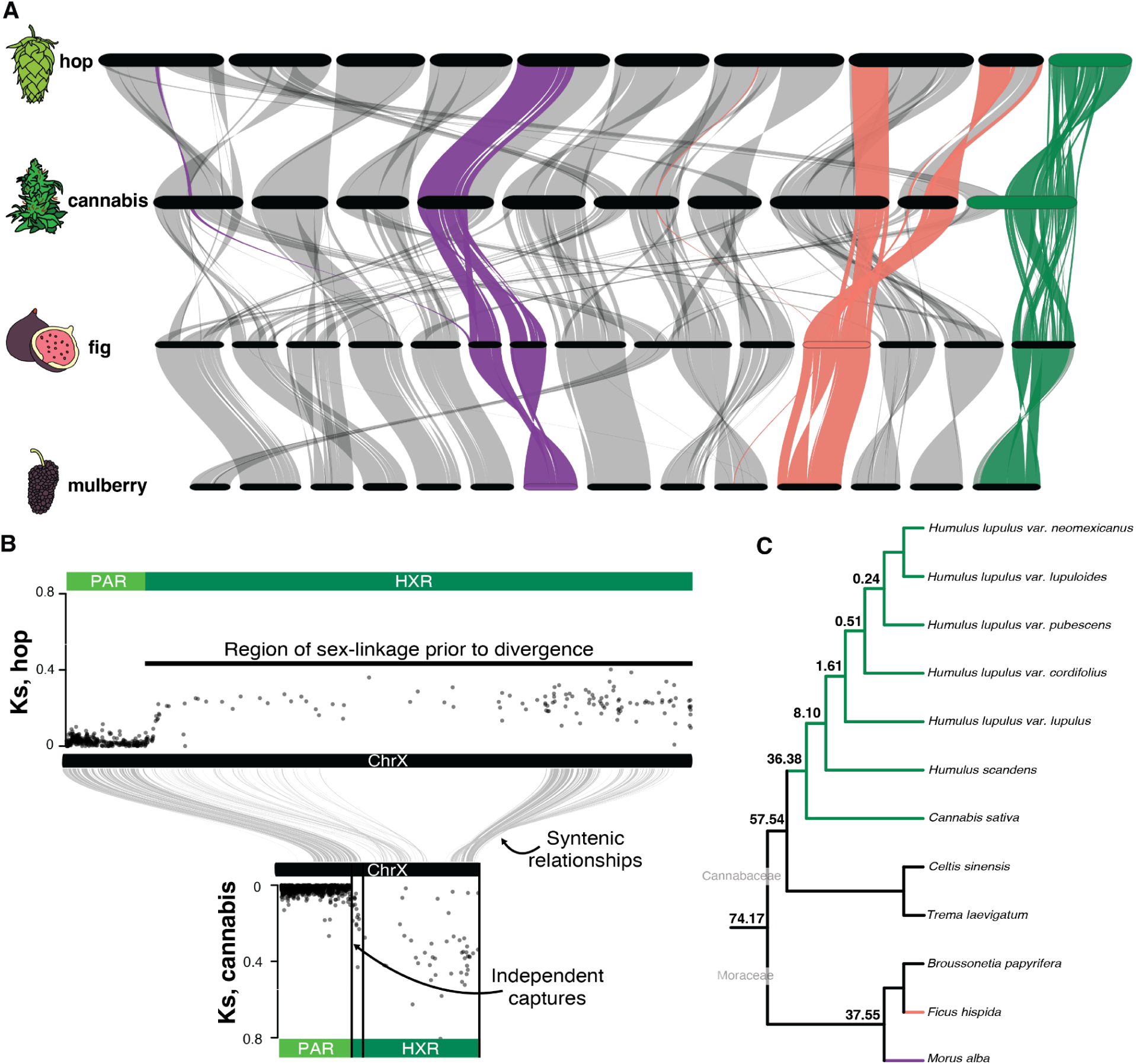
Identifying the origin of the sex chromosomes. **A)** GENESPACE synteny plot between hop, cannabis, fig, and mulberry. Green highlights syntenic relationships from the X chromosomes in hop and cannabis, while purple and salmon highlight synteny from the sex chromosomes of mulberry and fig, respectively. **B)** Synonymous protein changes (Ks) for one-to-one XY gametologs that highlight the homologous region of the X (HXR; values above one were removed). Syntenic relationships between the hop and cannabis X chromosomes were identified using GENESPACE. The hop reference is shown in its reverse complement, so that the HXR has the same orientation. **C)** Cladogram showing dates as the mean age in millions of years for species across Rosales.

Genes captured into the MSY in the same event are expected to have similar levels of Ks, where the older captures will have higher Ks between gametologs compared to younger captures (i.e., evolutionary strata (*37*)). While the max Ks in cannabis (0.81) is twice as high as hop (0.4), Ks is largely consistent when plotted across the X in both genera (Fig. 2), and we observed no evidence for a change point within this region that would indicate strata (Fig. S7), suggesting the vast majority of the MSY was captured in one event. In addition to the region of shared capture into the MSY, there is evidence for continual, but independent, gene captures in *Cannabis* and *Humulus*. In hop, we found evidence for nearly 50 Y-linked genes that were captured in the American hop, that are clearly in the PAR in *H. lupulus* var. *lupulus* (Fig. S5). In cannabis, Ks for gametologs along the MSY shows a clear, continuous pattern of addition from the PAR boundary at 29.3 Mb to 35.7 Mb, suggesting that genes in this region have been gradually added into the MSY over time, rather than through a large structural variant. Thus we found the Cannabaceae sex chromosomes have two regions: a region of independent captures and a shared region that was captured prior to the divergence of the two genera that likely contains the sex-determining gene.

### Y degeneration and faster-X evolution

The sex chromosomes have unique inheritance patterns that, coupled with the suppressed recombination of the MSY, can drive divergent molecular evolutionary dynamics. A classic observation in the MSY of sex chromosomes is degeneration. The suppressed recombination of the MSY reduces the efficacy of natural selection and can drive the accumulation of slightly deleterious mutations (*38*). As a consequence, the MSY tends to accumulate repetitive sequences, which can quickly increase the size of the Y, and over greater time can lead to gene loss and reduced size (*39*, *40*). If this linear trajectory holds true, then the extent of heteromorphy between XY pairs should be reflective of time. The Cannabaceae sex chromosomes defy these expectations as they have a shared origin but extreme cytological variation (Fig. 1), supporting the alternative hypothesis that time alone does not explain these changes (*11*). Like the Y, the X chromosome has unusual inheritance patterns. The X spends more evolutionary time in females, but the hemizygous state of the X chromosome in males is predicted to cause elevated rates of molecular evolution, a phenomenon referred to as the faster-X effect, that has mostly been investigated in animal systems thus far (*41*) (but see (*42*)).

The Cannabaceae sex chromosomes show clear signs of degeneration. In cannabis, which has the largest Y relative to its X counterpart, there is evidence for substantial gene loss (MSY models ∼46% of the homologous region of the X (HXR)). In *H. lupulus* var. *lupulus*, the MSY contains ∼17% the number in the HXR, with similarly low proportions in the Y chromosomes of vars. *lupuloides* and *neomexicanus* (both ∼20%; Table S5). Given the variation in the physical sizes of the Y chromosomes, one potential cause is through the differential loss of genes within the MSY. However, in hop, which shows the most extreme variation in size within a species, the number of XY gametologs are nearly the same, suggesting that the difference is driven by transposon accumulation. To test this, we annotated Transposable Elements (TE) using EDTA (*43*) and found the cannabis and hop genomes to be on average 74.3% and 86.8% repetitive, respectively (Table S4). TE content sharply increases at the beginning of the MSY (Fig. 1) and is 16% and 7% greater compared with the genome-wide averages for cannabis and hop, respectively (Table S6). Recent bursts in TE activity are evident on the Y chromosomes compared to their X counterparts, consistent with the increased overall percentages (Fig. 3). Importantly, the *H. lupulus* var. *lupulus* Y shows reduced TE expansion relative to the other Y chromosomes since the sex chromosomes evolved, which could explain its smaller size. Together these results support that the MSYs are highly-degenerated and that the cytological variation in Y chromosome sizes is a complex, ongoing interplay between gene loss and repeat expansion.

**Fig. 3.**
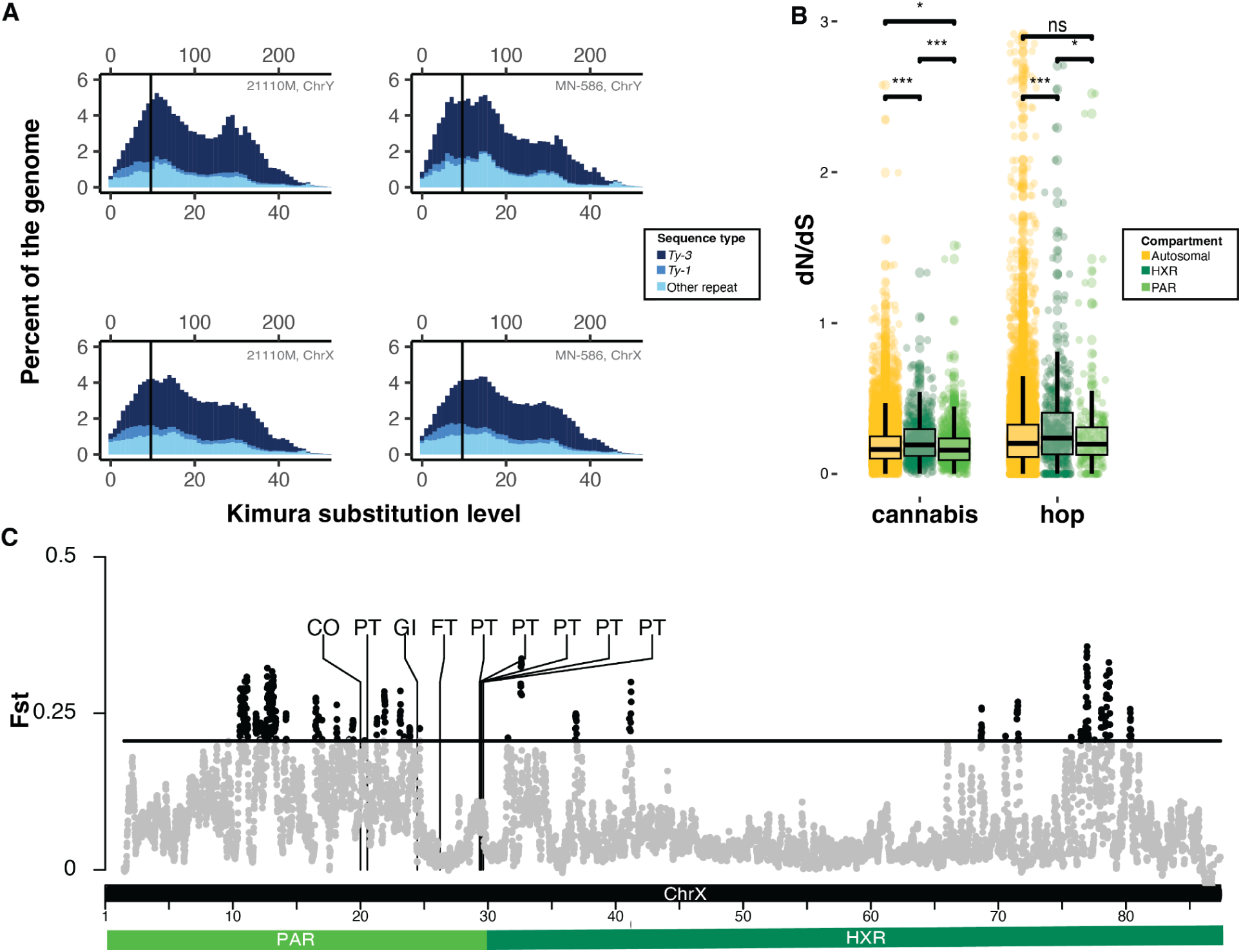
Molecular evolution of cannabis and hop XY chromosomes. **A**) Kimura substitution plots showing the repeat landscapes of the sex chromosomes. Values closer to 0 represent more recent events and higher values approaching 50 represent older events. Time was approximated by dividing the Kimura Distance by twice the mutation rate (using a value of 1x10^-9^), and the black lines indicate the time of the initial sex chromosome evolution. **B**) Boxplot of the ratio of nonsynonymous to synonymous protein changes (dN/dS). Asterisks indicate the level of significance from a Wilcoxon rank sum test with a Benjamini Hochberg correction for multiple tests. **C)** Fst outliers between chemotypes I-II vs III-IV in cannabis within 1Mb windows and a 100kb jump. Outliers were defined as values that were three standard deviations above the mean (an Fst of 20.6), and are indicated with black points.

The X chromosomes in cannabis and hop also show interesting patterns of molecular evolution. We used codeml in PAML (*44*) to calculate dN/dS ratios on 10,056 genes genome-wide. In cannabis, the HXR (N=597) has significantly higher dN/dS than the autosomes (N=8,671; Wilcoxon Rank Sum Test with Benjamini and Hochberg correction, p=8.3^-08) and PAR (N=782; p=8.2^-08) and in hop, we find the same pattern with the HXR (N=497) showing significantly greater dN/dS than the autosomes (N=7,932; p=0.00018) and PAR (N=226; p=0.02; Fig. 3). This supports that in both genera the X chromosomes show signatures of the faster-X effect. These molecular patterns are likely contributing to observations seen in some crosses, as there is ongoing evidence that sex chromosomes play a critical role in speciation (*45*).

### Key floral genes on the sex chromosomes

Flowering time is one of the several critical differences in the development of the sexes in cannabis and hop, wherein males flower earlier than females (*12*, *46*). The overall inflorescences are also architecturally different between the sexes. The male inflorescences consist of clusters of flowers (cymose panicles), while the female inflorescences are compound racemes (*12*). Within the female inflorescences are the glandular trichomes that produce the majority of the secondary metabolites, many of which are prenylated compounds (e.g., bitter acids and cannabinoids). It is currently unknown how much the sex chromosomes contribute to these sexually dimorphic traits.

Understanding flowering time is a major goal of many breeding programs in order to develop cultivars suitable for different latitudes and to control the timing of flowering in making controlled crosses. In cannabis, the appearance of male flowering typically occurs after producing four true leaf pairs (L4), 10-14 days prior to the appearance of female inflorescences (at nine true leaf pairs, L9; (*47*)). The photoperiod-dependent pathway for flowering includes genes such as *FLOWERING LOCUS T* (*FT*), *CONSTANS* (*CO*), and *GIGANTEA* (*GI*), and their conservation across plants could suggest similar functions in Cannabaceae (*48*, *49*). We found homologs for these three genes on the XYs of cannabis and hop (Table S7-S8). Consistent with these findings, GWAS and QTL analyses in cannabis identified peaks for flowering time across many loci, including the X chromosome (*50*). *FT* in particular stands out as a potential candidate for the difference in flowering time between the sexes, because in addition to the PAR copy and two autosomal homologs, there are copies that are found within the MSY and HXR. However, two observations might rule out *FT*. First, the origin of the duplicates to the MSY are recent and do not predate the divergence with hop, so seem unlikely to be driving the sex-specific differences in flowering given males also flower earlier in hop (Fig. S8). The second observation is related to gene expression of *FT*, which is expressed in leaf phloem cells and transported to the shoot apical meristem to initiate floral transition (*51*). Using RNAseq data of leaf tissue before and after flowering (*52*), the PAR *FT* is significantly differentially expressed, as well as the X-linked homolog in the HXR and autosomal homolog on Chr07 (Table S9). But none of the *FT* genes are significant when contrasting between the sexes at flowering. Another strong candidate is *FLOWERING LOCUS D* (*FD*), which interacts with *FT* to promote flowering (*53*). Gene tree topology supports that *FD* became sex-linked prior to the split between cannabis and hop (Fig. S8). Using RNAseq of apical meristems during flowering (L9 in female, L4 in male) (*54*), where *FD* is primarily expressed, the X copy is not significantly different between the sexes (adjusted p = 0.53). However, male expression is ten times higher on the Y-linked ortholog (Table S9). These results could suggest that *FD* is involved with the earlier timing of floral initiation found in male plants. Many additional floral regulators are located on the XYs, and a fine-stage developmental time series of gene expression, coupled with functional analyses, will help to better disentangle the roles of these genes.

The female inflorescences of hop and cannabis are economically important, since they produce high densities of glandular trichomes. In hop, the trichomes (called lupulin glands) produce the bitter acids, commonly referred to as alpha and beta acid, and essential oils used in brewing beer. Similarly, in cannabis, the female inflorescences produce the majority of the cannabinoids, such as cannabidiol (CBD) and tetrahydrocannabinol (THC). A critical step in the bitter acid and cannabinoid pathways is prenylation via aromatic prenyltransferases (*PT*) (*55*, *56*). In hop, two *PT* genes have been identified and are a tandem duplication located in the PAR of the XY chromosomes. The hop *PT1* has three to six homologs in cannabis, and several other *PT* genes are found in cannabis that vary both in function and copy number. Importantly, the *PT* in cannabis are also located in the PAR (*57*). Bitter acids and cannabinoids are major targets of breeding programs, so we tested for signs of selection among *PT* genes. We first used an Fst outlier approach using HiFi data for 49 female genotypes (*58*, *59*), where we contrasted between cannabis chemotypes I-II (higher THC; N=30) to III-IV (higher CBD; N=19). There is a local peak of higher Fst, which could in part be driven by fiber hemp samples in the chemotype III-IV class being bred for lower cannabinoid production. While *Cannabis* is monotypic, *Humulus* contains two species, which gives us the power to test for selection using the McDonald Kreitman test (MKT). Using WGS for 23 hops, we found *PT1* and *PT2* show signatures of adaptive variation (alpha = 0.76 and 0.1, respectively), of which *PT1* shows the strongest signal under Fisher’s exact test (p=0.03, FDR= n.s.). Given that *PT* genes prenylate CBD, THC, and the bitter acids, robust comparisons that contrast the abundance of metabolite production may be more revealing.

A curious pattern is evident when contrasting cannabis and hop for these genes. Cannabis has evolved many more copies of *PT*, with diverse functions, and shows copy-number variation among genotypes, while hop is static. Moreover, the gradual expansion of the MSY in cannabis leaves the closest *PT* gene ∼1.1Mb away from the boundary. This gradual expansion of the MSY is potentially related to centromere proximity (Fig. S9) as the existing reduced recombination could facilitate sex-linkage (*60*, *61*). However, given that females are strongly selected in cannabis, instead the MSY boundary may have expanded due to this sexually antagonistic selection (*27*, *62*). Indeed, recent analyses show the boundary of the MSY is variable in cannabis, with some genotypes supporting this boundary expanding further into the PAR (*58*), closer to the *PT* and *FT* genes. While altogether these results suggest the sex chromosomes played a role in the domestication of these species, it is also possible that the speciation process and subsequent adaptation has also helped to shape the sex chromosomes.

### Evidence for a background, not active, Y system

It is well established that ethylene plays a demonstrable role in sex determination in cannabis and hop. Ethephon treatment when applied to XY plants, which releases ethylene, will initiate female inflorescence development, rather than male (*30*, *63*). The opposite pattern has also been demonstrated, wherein ethylene inhibitors promote male development on XX plants (*64*). These observations suggest that genes that determine sex are involved with the ethylene pathway. The sex determination genes are also expected to act early in flower development, because the dimorphic architectures between the sexes are apparent shortly after floral meristem initiation (*12*) and neither sex produces vestigial structures of the other sex (*47*). Analyses to determine the genetic basis for the sex-determining genes using trait-based approaches and transcriptomics (*54*, *65*) have made promising discoveries, but so far no clear candidate has been identified.

Given the shared origin of a sex chromosome, we searched for highly-conserved genes in the MSYs. Sixty three genes are shared on the Y chromosomes between the genera, 13 of which are suspected to be involved with floral development based on orthologous functions (Table S8). None of these appear to be obvious sex-determination genes based on their developmental timing and function. For instance, the maintenance of SIGNAL PEPTIDE PEPTIDASE (*AtSPP*) and ROP ENHANCER 1 (*REN1*), which in *Arabidopsis* are involved in pollen germination (*66*) and pollen tube growth (*67*), respectively, suggests a role in male fertility in Cannabaceae, but not overall stamen development. Moreover, when contrasting male gene expression at the L2 stage (pre-flowering) to the L4 stage (flowering), only three MSY genes were significantly differentially expressed, but none of these are shared with hop nor have clear roles in reproductive organ development (Table S10). The lack of an obvious sex-determining gene on the Y tracks with crossing experiments using polyploids that vary in X copy number. In an active-Y sex determination system, inheritance of the Y is expected to lead to male individuals, regardless of the number of X copies (*68*). In hop and cannabis diploids, XY individuals typically develop strictly male inflorescences, whereas polyploid individuals with XXY or XXXY karyotypes commonly produce female flowers or are monoecious (*69*, *70*). These lines of evidence suggest that the sex-determining gene instead resides on the X. The maintained pollen and flowering timing genes, like *FD*, in the MSY suggest that the Y is not inactive, but plays other critical fertility roles, which we term a “background-Y”.

### A candidate X-linked sex-determination gene

Natural variation in sexual systems can be used to overcome limitations in functional genomics in order to identify sex-determining genes (*71*). In cannabis, reversions to monoecy, where a plant can produce both functional female and male flowers, have been observed in XX and XY diploids. One possible route to transitioning from dioecy to monoecy in an XX individual involves movement of sequence from the MSY to the X chromosome. To test this hypothesis, we examined the genomes for the monoecious ‘Futura 75’ and ‘Uso 31’ genotypes. There are no signatures of the Y chromosome on the X_m_, including no dense peaks of Y-mers ( Fig. 1; Table S2), suggesting they are both XX in their karyotype, and assessing 1,096 gene trees showed no evidence of a topology suggesting an MSY gene translocated. These results suggest that in these individuals, monoecy evolved through a modification of a gene on the X chromosome, likely within the sex-determining gene, consistent with the background-Y hypothesis.

To uncover the genetic basis for monoecy in cannabis, we generated a reciprocal backcross (BC_1_) between a female and monoecious line and genotyped 950 BC_1_ offspring with low-pass Illumina shotgun sequencing. We mapped reads to a pangenome graph using the two male and two monecious cannabis references and used KhufuPAN (*72*) to identify loci associated with monoecy. All 19 hits are located on the X chromosome, with 73.7% of these landing between 77.6-81.7 Mb (Fig. 4). We generated PacBio HiFi data for 10 additional monoecious genotypes and used existing HiFi reads from 51 cannabis genotypes (*58*, *59*) outside of the BC_1_ and also ran an F_st_ outlier analysis (N=49 female, N=14 monoecious). 66.8% of the outliers are found on the X chromosome, and particularly surrounding the locus identified with the BC_1_ (Fig. 4), strongly suggesting this is the locus associated with the monoecious trait.

**Fig. 4.**
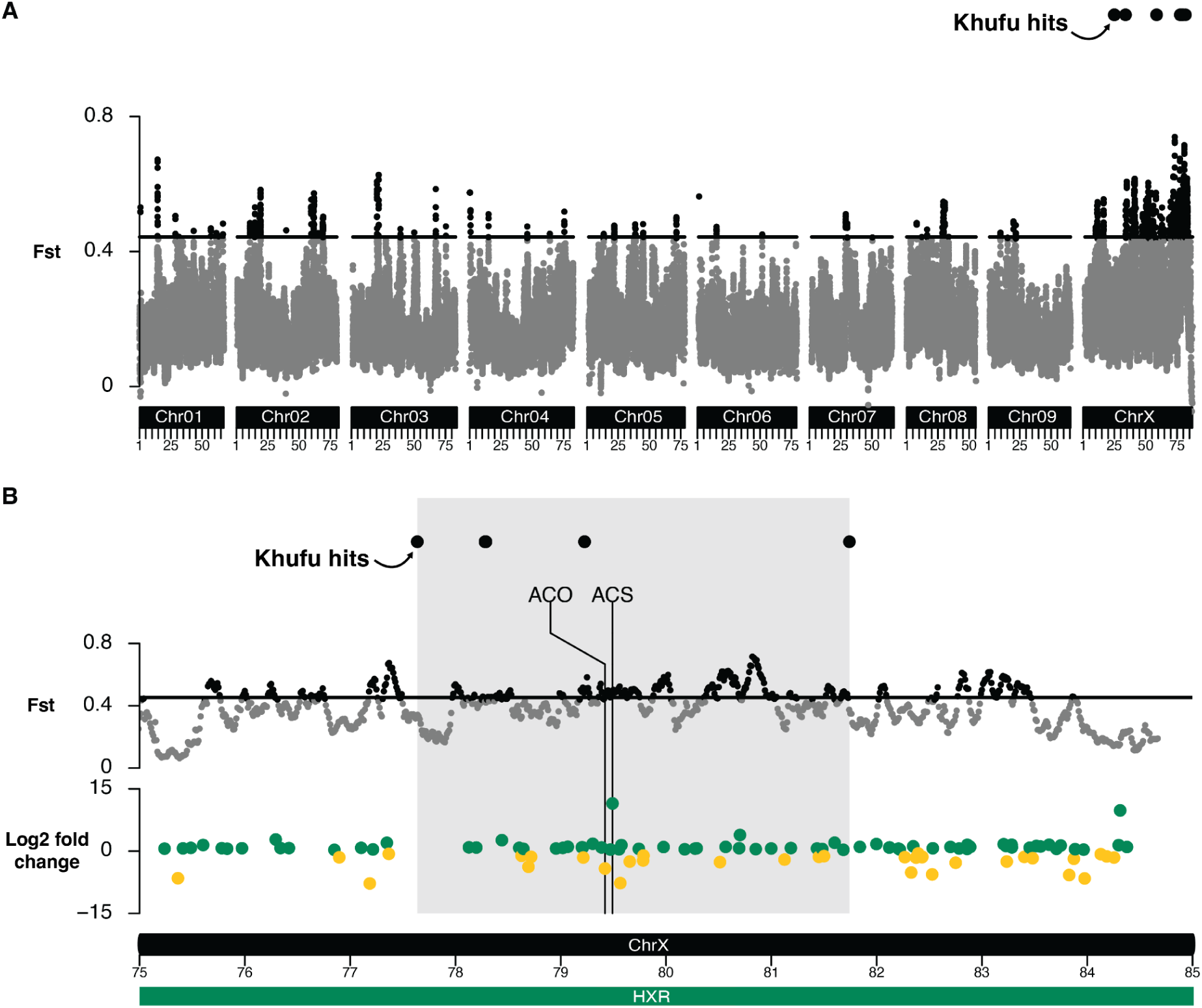
Identifying the sex-determining locus in cannabis. **A)** karyoplot showing Khufu hits from a backcross segregating for monoecy and Fst outliers between female and monoecious cannabis genotypes. Fst was calculed within 1Mb windows and a 100kb jump. Outliers were defined as values that were three standard deviations above the mean (an Fst of 43.8), and are indicated with black points. **B)** the region identified as the monoecy locus is highlighted in light gray, with the same Khufu hits and Fst outliers as in panel A. Also shown are the location of genes that are significantly differentially expressed between the sexes in cannabis during flowering (adjusted p-value < 0.05). The positive axis are genes with greater expression in females, the negative axis are greater in males. Homologs of two ethylene biosynthesis genes are located in this locus, ACC synthase (*ACS*) and ACC oxidase (*ACO*).

We next analyzed gene expression between the sexes during flowering, to further narrow in on candidate sex-determining genes. Forty six genes were significantly differentially expressed in the monoecy region; 11 are related to flowering or ethylene (Table S11). None of the floral genes are obvious candidates for sex-determination; however, because of the demonstrated role of ethylene in promoting female inflorescence development, the ACO and ACS genes are notable candidates. The ethylene biosynthesis pathway involves two enzymatic steps. First, S-adenosyl-L-methionine is converted into 1-aminocyclopropane-1-carboxylic acid (ACC) by ACC synthase (*ACS*), then ACC is converted to ethylene by ACC oxidase (*ACO*) (*73–77*). The *ACO* gene located in the monoecy locus is orthologous to *ACO5* (Fig S8) and expression of this gene was greater in males, which is counter to the expectation that female flowers require ethylene (Table S11). Moreover, the most abundant tissue that it is expressed in are roots in cannabis (Fig. S10). The *ACS* gene, however, is in the clade with other type II ACS genes, and was exclusively expressed in females and monoecious plants within floral tissues and stems during flowering (Fig S8, S11), which are patterns consistent with the ACS genes that have been demonstrated to determine sex in cucurbits (*78*, *79*).

In some crosses, it has been shown that monoecy is a recessive trait (*6*, *80*). We searched for variants that were homozygous in monoecious genotypes that could be causative. We used a combination of Sawfish, DeepVariant, and mpileup to call SNPs, INDELS, and structural variants (SV). No large SVs were identified within the *ACO5* and *ACS* gene models, and no SNP or INDEL segregated in *ACO5* for monoecy. However, we found two INDELS, each three basepairs in length, in exon 4 of *ACS* just outside an aminotransferase domain, that are homozygous in 13/14 monoecious genotypes (one ‘Futura 75’ was heterozygous, but another ‘Futura 75’ was homozygous). Of the 49 female genotypes scanned, only two had the INDELs, but were both heterozygous. These results lead us to the striking conclusion that sex is determined in cannabis and hop using the same ethylene-based sex determination pathway as in cucurbits, which are more than 100 million years diverged.

## Conclusions

Understanding the genetic basis for sex has been a topic of interest for over a century. Cytologists, like Nettie Stevens, uncovered the inheritance of the heteromorphic sex chromosome that determined maleness (*8*). While both cannabis and hop were among the first plants to have their sex chromosomes identified a century ago due to their heteromorphic XYs, uncovering the genes that determine sex has remained elusive. Our near-complete genome assemblies have shown the evolution of sex chromosomes in cannabis and hop in unprecedented detail. The ancient MSY contains male-fertility genes, but given the lack of clear genes associated with sex determination, support that it is a background-Y rather than an active-Y. The X, however, has demonstrably been shown to be involved with sex determination through crosses and the reversion to monoecy. Here we identified the gene that promotes female inflorescence development is the ethylene pathway gene *ACS*. While some of these results are in line with patterns found in mammals, like gene loss in the MSY and faster-X, an open question remains on how background-Y and an X chromosome that contain the sex determining gene may evolve differently. Altogether, the diverse sex chromosomes, and their associated reproductive genes in both cannabis and hop, played a pivotal role in the domestication, breeding, and ongoing production of these species.

## Acknowledgments

We would like to thank the US Department of Agriculture Hemp Germplasm Laboratory (8060-21000-034-000-D) for generously providing materials for this manuscript. We would also like to thank the HudsonAlpha IT Department for their research computing support. **Funding:** Support for this work was provided by the US Department of Agriculture National Institute of Food and Agriculture Postdoctoral Fellowship (USDA NIFA) no. 2022-67012-38987 (S.B.C.), USDA NIFA no. 2023-67013-39620 (A.H.), National Science Foundation (NSF) IOS-PGRP CAREER no. 2239530 (A.H.), USDA Non-Assistance Cooperative Agreement (NACA) no. 58-8060-4-002 (A.H., Z.S.), NIH R35 GM147107 (R.F.G.), and Institute of *Cannabis* Research at Colorado State University Pueblo ICR-FY22-Kane (R.F.G., D.V., N.C.K.).

## Author contributions

Conceptualization: S.B.C., J.S.H., J.K.M., G.J.M., A.H. Funding acquisition: S.B.C., Z.S., R.F.G., D.V., N.C.K., A.H. Formal analysis: S.B.C., P.C.B., Z.A.M., W.K., L.P.-C., R.C.L., N.A., A.O. Investigation: S.B.C., J.M., Z.S., G.S., T.G., R.F., H.H., H.M., Z.M., J.G., J.K.M. Methodology: S.B.C, A.H. Resources: J.S.H., Z.S., G.S., T.G., K.A.E., Z.E.M., L.R.O., L.B.S., N.C.K., G.J.M., J.C., A.H. Visualization: S.B.C., P.C.B., J.T.L., L.M.A., Z.A.M. Writing - original draft: S.B.C., A.H. Writing - review and editing: S.B.C., P.C.B., J.T.L., L.M.A., J.S.H., L.P.-C., R.C.L., Z.S., G.S., K.A.E., L.B.S., D.V., R.F.G., N.C.K., J.K.M., T.P.M., A.H.

## Competing interests

A.H. and J.C. are co-founders and board members of Veil Genomics, a long-read genotyping company. D.V. is a board member of the 501(C)3 non-profit Agricultural Genomics Foundation and the sole owner of the company CGRI, LLC. J.K.M. and R.F. are founders of New West Genetics, a breeding program that produces hybrid hemp seed. **Data and materials availability:** The genome assemblies have been deposited on NCBI under BioProjects (TBD). Sequencing libraries for the genome assembly and annotation are publicly available on NCBI under BioProject (TBD). The remaining sequencing generated in support of this manuscript can be found under BioProject (TBD). Scripts that support this manuscript can be found at https://github.com/sarahcarey/Cannabaceae_sexChromosome. Additional files can be found on Zenodo 10.5281/zenodo.16326991.

## Supplementary Materials

Materials and Methods

Supplementary Text

Figs. S1 to S12

Tables S1 to S16

References (81-152)

## Notes

### Summary of Updates

This manuscript has been revised to include analyses to uncover a candidate sex determining gene

## References

1. G. Ren, X. Zhang, Y. Li, K. Ridout, M. L. Serrano-Serrano, Y. Yang, A. Liu, G. Ravikanth, M. A. Nawaz, A. S. Mumtaz, N. Salamin, L. Fumagalli, Large-scale whole-genome resequencing unravels the domestication history of Cannabis sativa. Sci. Adv. 7 (2021).

2. E. Small, D. Marcus, Hemp: A new crop with new uses for North America. Trends in new crops and new uses 24, 284–326 (2002).

3. I. S. Hornsey, Brewing (Royal Society of Chemistry, 2013).

4. J. Harrison, EFFECT OF HOP SEEDS ON BEER QUALITY. J. Inst. Brew. 77, 350–352 (1971).

5. C. Lipson Feder, O. Cohen, A. Shapira, I. Katzir, R. Peer, O. Guberman, S. Procaccia, P. Berman, M. Flaishman, D. Meiri, Fertilization following pollination predominantly decreases phytocannabinoids accumulation and alters the accumulation of terpenoids in Cannabis inflorescences. Front. Plant Sci. 12, 753847 (2021).

6. O. V. Razumova, O. S. Alexandrov, M. G. Divashuk, T. I. Sukhorada, G. I. Karlov, Molecular cytogenetic analysis of monoecious hemp (Cannabis sativa L.) cultivars reveals its karyotype variations and sex chromosomes constitution. Protoplasma 253, 895–901 (2016).

7. L. Garcia-de Heer, J. Mieog, A. Burn, T. Kretzschmar, Why not XY? Male monoecious sexual phenotypes challenge the female monoecious paradigm in Cannabis sativa L. Front. Plant Sci. 15, 1412079 (2024).

8. N. M. Stevens, Studies in Spermatogenesis .. (1905).

9. O. Winge, On sex chromosomes, sex determination and preponderance of females in some dioecious plants. Compt. Rend. Trav. Lab. Carlsberg 15, 1–26 (1923).

10. K. Hirata, Sex reversal in hemp. Jour. Soc. Agr. For. Sapporo (1924).

11. S. S. Renner, N. A. Müller, Plant sex chromosomes defy evolutionary models of expanding recombination suppression and genetic degeneration. Nat Plants 7, 392–402 (2021).

12. H. L. Shephard, J. S. Parker, P. Darby, C. C. Ainsworth, Sexual development and sex chromosomes in hop. New Phytol. 148, 397–411 (2000).

13. D. Prentout, N. Stajner, A. Cerenak, T. Tricou, C. Brochier-Armanet, J. Jakse, J. Käfer, G. A. B. Marais, Plant genera Cannabis and Humulus share the same pair of well-differentiated sex chromosomes. New Phytol. 231, 1599–1611 (2021).

14. A. H. Sinclair, P. Berta, M. S. Palmer, J. Ross Hawkins, B. L. Griffiths, M. J. Smith, J. W. Foster, A. Frischauf, R. Lovell-Badge, P. N. Goodfellow, A gene from the human sex-determining region encodes a protein with homology to a conserved DNA-binding motif. Nature 346, 240–244 (1990).

15. J. S. Parker, M. S. Clark, Dosage sex-chromosome systems in plants. Plant Sci. 80, 79–92 (1991).

16. S. Gao, B. Wang, S. Xie, X. Xu, J. Zhang, L. Pei, Y. Yu, W. Yang, Y. Zhang, A high-quality reference genome of wild Cannabis sativa. Hortic Res 7, 73 (2020).

17. L. K. Padgitt-Cobb, N. J. Pitra, P. D. Matthews, J. A. Henning, D. A. Hendrix, An improved assembly of the “Cascade” hop (Humulus lupulus) genome uncovers signatures of molecular evolution and refines time of divergence estimates for the Cannabaceae family. Horticulture Research 10, uhac281 (2023).

18. A. Rhie, S. A. McCarthy, O. Fedrigo, J. Damas, G. Formenti, S. Koren, M. Uliano-Silva, W. Chow, A. Fungtammasan, J. Kim, C. Lee, B. J. Ko, M. Chaisson, G. L. Gedman, L. J. Cantin, F. Thibaud-Nissen, L. Haggerty, I. Bista, M. Smith, B. Haase, J. Mountcastle, S. Winkler, S. Paez, J. Howard, S. C. Vernes, T. M. Lama, F. Grutzner, W. C. Warren, C. N. Balakrishnan, D. Burt, J. M. George, M. T. Biegler, D. Iorns, A. Digby, D. Eason, B. Robertson, T. Edwards, M. Wilkinson, G. Turner, A. Meyer, A. F. Kautt, P. Franchini, H. W. Detrich 3rd, H. Svardal, M. Wagner, G. J. P. Naylor, M. Pippel, M. Malinsky, M. Mooney, M. Simbirsky, B. T. Hannigan, T. Pesout, M. Houck, A. Misuraca, S. B. Kingan, R. Hall, Z. Kronenberg, I. Sović, C. Dunn, Z. Ning, A. Hastie, J. Lee, S. Selvaraj, R. E. Green, N. H. Putnam, I. Gut, J. Ghurye, E. Garrison, Y. Sims, J. Collins, S. Pelan, J. Torrance, A. Tracey, J. Wood, R. E. Dagnew, D. Guan, S. E. London, D. F. Clayton, C. V. Mello, S. R. Friedrich, P. V. Lovell, E. Osipova, F. O. Al-Ajli, S. Secomandi, H. Kim, C. Theofanopoulou, M. Hiller, Y. Zhou, R. S. Harris, K. D. Makova, P. Medvedev, J. Hoffman, P. Masterson, K. Clark, F. Martin, K. Howe, P. Flicek, B. P. Walenz, W. Kwak, H. Clawson, M. Diekhans, L. Nassar, B. Paten, R. H. S. Kraus, A. J. Crawford, M. T. P. Gilbert, G. Zhang, B. Venkatesh, R. W. Murphy, K.-P. Koepfli, B. Shapiro, W. E. Johnson, F. Di Palma, T. Marques-Bonet, E. C. Teeling, T. Warnow, J. M. Graves, O. A. Ryder, D. Haussler, S. J. O’Brien, J. Korlach, H. A. Lewin, K. Howe, E. W. Myers, R. Durbin, A. M. Phillippy, E. D. Jarvis, Towards complete and error-free genome assemblies of all vertebrate species. Nature 592, 737–746 (2021).

19. A. Rhie, S. Nurk, M. Cechova, S. J. Hoyt, D. J. Taylor, N. Altemose, P. W. Hook, S. Koren, M. Rautiainen, I. A. Alexandrov, J. Allen, M. Asri, A. V. Bzikadze, N.-C. Chen, C.-S. Chin, M. Diekhans, P. Flicek, G. Formenti, A. Fungtammasan, C. Garcia Giron, E. Garrison, A. Gershman, J. L. Gerton, P. G. S. Grady, A. Guarracino, L. Haggerty, R. Halabian, N. F. Hansen, R. Harris, G. A. Hartley, W. T. Harvey, M. Haukness, J. Heinz, T. Hourlier, R. M. Hubley, S. E. Hunt, S. Hwang, M. Jain, R. K. Kesharwani, A. P. Lewis, H. Li, G. A. Logsdon, J. K. Lucas, W. Makalowski, C. Markovic, F. J. Martin, A. M. Mc Cartney, R. C. McCoy, J. McDaniel, B. M. McNulty, P. Medvedev, A. Mikheenko, K. M. Munson, T. D. Murphy, H. E. Olsen, N. D. Olson, L. F. Paulin, D. Porubsky, T. Potapova, F. Ryabov, S. L. Salzberg, M. E. G. Sauria, F. J. Sedlazeck, K. Shafin, V. A. Shepelev, A. Shumate, J. M. Storer, L. Surapaneni, A. M. Taravella Oill, F. Thibaud-Nissen, W. Timp, M. Tomaszkiewicz, M. R. Vollger, B. P. Walenz, A. C. Watwood, M. H. Weissensteiner, A. M. Wenger, M. A. Wilson, S. Zarate, Y. Zhu, J. M. Zook, E. E. Eichler, R. J. O’Neill, M. C. Schatz, K. H. Miga, K. D. Makova, A. M. Phillippy, The complete sequence of a human Y chromosome. Nature 621, 344–354 (2023).

20. A. Muyle, J. Käfer, N. Zemp, S. Mousset, F. Picard, G. A. Marais, SEX-DETector: A probabilistic approach to study sex chromosomes in non-model organisms. Genome Biol. Evol. 8, 2530–2543 (2016).

21. J. Käfer, N. Lartillot, G. A. B. Marais, F. Picard, Detecting sex-linked genes using genotyped individuals sampled in natural populations. Genetics 218 (2021).

22. S. B. Carey, J. T. Lovell, J. Jenkins, J. Leebens-Mack, J. Schmutz, M. A. Wilson, A. Harkess, Representing sex chromosomes in genome assemblies. Cell Genom. 2, 100132 (2022).

23. S. B. Carey, L. Aközbek, J. T. Lovell, J. Jenkins, A. L. Healey, S. Shu, P. Grabowski, A. Yocca, A. Stewart, T. Jones, K. Barry, S. Rajasekar, J. Talag, C. Scutt, P. P. Lowry 2nd, J. Munzinger, E. B. Knox, D. E. Soltis, P. S. Soltis, J. Grimwood, J. Schmutz, J. Leebens-Mack, A. Harkess, ZW sex chromosome structure in Amborella trichopoda. Nat. Plants, doi: 10.1038/s41477-024-01858-x (2024).

24. A. Haunold, Hop Production, Breeding, and Variety Development in Various Countries. J. Am. Soc. Brew. Chem. 39, 27–34 (1981).

25. H. Cheng, G. T. Concepcion, X. Feng, H. Zhang, H. Li, Haplotype-resolved de novo assembly using phased assembly graphs with hifiasm. Nat. Methods 18, 170–175 (2021).

26. A. Rhie, B. P. Walenz, S. Koren, A. M. Phillippy, Merqury: reference-free quality, completeness, and phasing assessment for genome assemblies. Genome Biol. 21, 245 (2020).

27. S. P. Otto, J. R. Pannell, C. L. Peichel, T.-L. Ashman, D. Charlesworth, A. K. Chippindale, L. F. Delph, R. F. Guerrero, S. V. Scarpino, B. F. McAllister, About PAR: the distinct evolutionary dynamics of the pseudoautosomal region. Trends Genet. 27, 358–367 (2011).

28. Q. Zhang, R. E. Onstein, S. A. Little, H. Sauquet, Estimating divergence times and ancestral breeding systems in Ficus and Moraceae. Ann. Bot. 123, 191–204 (2019).

29. T. Dennis Thomas, In Vitro Modification of Sex Expression in Mulberry (Morus Alba) by Ethrel and Silver Nitrate. Plant Cell Tissue Organ Cult. 77, 277–281 (2004).

30. H. Y. Mohan Ram, R. Sett, Modification of growth and sex expression in cannabis sativa by aminoethoxyvinylglycine and ethephon. Z. Pflanzenphysiol. 105, 165–172 (1982).

31. X. Zhang, G. Wang, S. Zhang, S. Chen, Y. Wang, P. Wen, X. Ma, Y. Shi, R. Qi, Y. Yang, Z. Liao, J. Lin, J. Lin, X. Xu, X. Chen, X. Xu, F. Deng, L. Zhao, Y.-L. Lee, R. Wang, X.-Y. Chen, Y.-R. Lin, J. Zhang, H. Tang, J. Chen, R. Ming, Genomes of the Banyan Tree and Pollinator Wasp Provide Insights into Fig-Wasp Coevolution. Cell 183, 875–889.e17 (2020).

32. Z. Xia, X. Dai, W. Fan, C. Liu, M. Zhang, P. Bian, Y. Zhou, L. Li, B. Zhu, S. Liu, Z. Li, X. Wang, M. Yu, Z. Xiang, Y. Jiang, A. Zhao, Chromosome-level Genomes Reveal the Genetic Basis of Descending Dysploidy and Sex Determination in Morus Plants. Genomics Proteomics Bioinformatics, doi: 10.1016/j.gpb.2022.08.005 (2022).

33. O. V. Razumova, M. G. Divashuk, O. S. Alexandrov, G. I. Karlov, GISH painting of the Y chromosomes suggests advanced phases of sex chromosome evolution in three dioecious Cannabaceae species (Humulus lupulus, H. japonicus, and Cannabis sativa). Protoplasma 260, 249–256 (2023).

34. K. J. Hoff, A. Lomsadze, M. Stanke, M. Borodovsky, BRAKER2: incorporating protein homology information into gene prediction with GeneMark-EP and AUGUSTUS. Plant and Animal Genomes XXVI (2018).

35. L. Gabriel, T. Brůna, K. J. Hoff, M. Ebel, A. Lomsadze, M. Borodovsky, M. Stanke, BRAKER3: Fully automated genome annotation using RNA-seq and protein evidence with GeneMark-ETP, AUGUSTUS, and TSEBRA. Genome Res., doi: 10.1101/gr.278090.123 (2024).

36. C. Zhang, M. Rabiee, E. Sayyari, S. Mirarab, ASTRAL-III: polynomial time species tree reconstruction from partially resolved gene trees. BMC Bioinformatics 19, 153 (2018).

37. B. T. Lahn, D. C. Page, Four evolutionary strata on the human X chromosome. Science 286, 964–967 (1999).

38. D. Charlesworth, B. Charlesworth, G. Marais, Steps in the evolution of heteromorphic sex chromosomes. Heredity 95, 118–128 (2005).

39. A. S. T. Papadopulos, M. Chester, K. Ridout, D. A. Filatov, Rapid Y degeneration and dosage compensation in plant sex chromosomes. Proc. Natl. Acad. Sci. U. S. A. 112, 13021–13026 (2015).

40. B. Sacchi, Z. Humphries, J. Kružlicová, M. Bodláková, C. Pyne, B. I. Choudhury, Y. Gong, V. Bačovský, R. Hobza, S. C. H. Barrett, S. I. Wright, Phased assembly of Neo-sex chromosomes reveals extensive Y degeneration and rapid genome evolution in Rumex hastatulus. Mol. Biol. Evol. 41, msae074 (2024).

41. J. E. Mank, B. Vicoso, S. Berlin, B. Charlesworth, Effective population size and the Faster-X effect: empirical results and their interpretation. Evolution 64, 663–674 (2010).

42. M. Krasovec, B. Nevado, D. A. Filatov, A comparison of selective pressures in plant X-linked and autosomal genes. Genes (Basel) 9, 234 (2018).

43. S. Ou, W. Su, Y. Liao, K. Chougule, J. R. A. Agda, A. J. Hellinga, C. S. B. Lugo, T. A. Elliott, D. Ware, T. Peterson, N. Jiang, C. N. Hirsch, M. B. Hufford, Benchmarking transposable element annotation methods for creation of a streamlined, comprehensive pipeline. Genome Biol. 20, 275 (2019).

44. Z. Yang, PAML 4: phylogenetic analysis by maximum likelihood. Mol. Biol. Evol. 24, 1586–1591 (2007).

45. B. A. Payseur, D. C. Presgraves, D. A. Filatov, Introduction: Sex chromosomes and speciation. Mol. Ecol. 27, 3745–3748 (2018).

46. V. Mediavilla, M. Jonquera, I. Schmid-Slembrouck, Decimal code for growth stages of hemp (Cannabis sativa L.). (1998).

47. J. Shi, S. Schilling, R. Melzer, Morphological and genetic analysis of inflorescence and flower development in hemp (*Cannabis sativa*L.), bioRxiv (2024). 10.1101/2024.01.25.577276.

48. C. Brandoli, C. Petri, M. Egea-Cortines, J. Weiss, Gigantea: Uncovering new functions in flower development. Genes (Basel) 11, 1142 (2020).

49. J. M. Romero, G. Serrano-Bueno, C. Camacho-Fernández, M. H. Vicente, M. T. Ruiz, J. R. Pérez-Castiñeira, J. Pérez-Hormaeche, F. T. S. Nogueira, F. Valverde, CONSTANS, a HUB for all seasons: How photoperiod pervades plant physiology regulatory circuits. Plant Cell 36, 2086–2102 (2024).

50. L. Steel, M. Welling, N. Ristevski, K. Johnson, A. Gendall, Comparative genomics of flowering behavior in Cannabis sativa. Front. Plant Sci. 14 (2023).

51. L. Corbesier, C. Vincent, S. Jang, F. Fornara, Q. Fan, I. Searle, A. Giakountis, S. Farrona, L. Gissot, C. Turnbull, G. Coupland, FT protein movement contributes to long-distance signaling in floral induction of Arabidopsis. Science 316, 1030–1033 (2007).

52. C. A. Dowling, T. P. Michael, P. F. McCabe, S. Schilling, R. Melzer, *FT*-like genes in Cannabis and hops: sex specific expression and copy-number variation may explain flowering time variation, bioRxiv (2024). 10.1101/2024.10.04.616617.

53. M. Abe, Y. Kobayashi, S. Yamamoto, Y. Daimon, A. Yamaguchi, Y. Ikeda, H. Ichinoki, M. Notaguchi, K. Goto, T. Araki, FD, a bZIP protein mediating signals from the floral pathway integrator FT at the shoot apex. Science 309, 1052–1056 (2005).

54. J. Shi, M. Toscani, C. A. Dowling, S. Schilling, R. Melzer, Identification of genes associated with sex expression and sex determination in hemp (Cannabis sativa L.). J. Exp. Bot., doi: 10.1093/jxb/erae429 (2024).

55. H. Li, Z. Ban, H. Qin, L. Ma, A. King, G. Wang, A heteromeric membrane-bound prenyltransferase complex from hop catalyzes three sequential aromatic prenylations in the bitter acid Pathway1[OPEN]. Plant Physiol. 167, 650–659 (2015).

56. X. Luo, M. A. Reiter, L. d’Espaux, J. Wong, C. M. Denby, A. Lechner, Y. Zhang, A. T. Grzybowski, S. Harth, W. Lin, H. Lee, C. Yu, J. Shin, K. Deng, V. T. Benites, G. Wang, E. E. K. Baidoo, Y. Chen, I. Dev, C. J. Petzold, J. D. Keasling, Complete biosynthesis of cannabinoids and their unnatural analogues in yeast. Nature 567, 123–126 (2019).

57. K. A. Rea, J. A. Casaretto, M. S. Al-Abdul-Wahid, A. Sukumaran, J. Geddes-McAlister, S. J. Rothstein, T. A. Akhtar, Biosynthesis of cannflavins A and B from Cannabis sativa L. Phytochemistry 164, 162–171 (2019).

58. R. C. Lynch, L. K. Padgitt-Cobb, A. R. Garfinkel, B. J. Knaus, N. T. Hartwick, N. Allsing, A. Aylward, P. C. Bentz, S. B. Carey, A. Mamerto, J. K. Kitony, K. Colt, E. R. Murray, T. Duong, H. I. Chen, A. Trippe, A. Harkess, S. Crawford, K. Vining, T. P. Michael, Domesticated cannabinoid synthases amid a wild mosaic cannabis pangenome. Nature, doi: 10.1038/s41586-025-09065-0 (2025).

59. G. M. Stack, M. A. Quade, D. G. Wilkerson, L. A. Monserrate, P. C. Bentz, S. B. Carey, J. Grimwood, J. A. Toth, S. Crawford, A. Harkess, L. B. Smart, Comparison of recombination rate, reference bias, and unique pangenomic haplotypes in Cannabis sativa using seven DE Novo genome assemblies. Int. J. Mol. Sci. 26 (2025).

60. Q. Yu, S. Hou, R. Hobza, F. A. Feltus, X. Wang, W. Jin, R. L. Skelton, A. Blas, C. Lemke, J. H. Saw, P. H. Moore, M. Alam, J. Jiang, A. H. Paterson, B. Vyskot, R. Ming, Chromosomal location and gene paucity of the male specific region on papaya Y chromosome. Mol. Genet. Genomics 278, 177–185 (2007).

61. H. She, Z. Liu, S. Li, Z. Xu, H. Zhang, F. Cheng, J. Wu, X. Wang, C. Deng, D. Charlesworth, W. Gao, W. Qian, Evolution of the spinach sex-linked region within a rarely recombining pericentromeric region. Plant Physiol. 193, 1263–1280 (2023).

62. W. Rice, Sex chromosomes and the evolution of sexual dimorphism. Evolution 38 (1984).

63. E. Galoch, The hormonal control of sex differentiation in dioecious plants of hemp (Cannabis sativd). The influence of plant growth regulators on sex expression in male and female plants. Acta Soc. Bot. Pol. Pol. Tow. Bot. 47, 153–162 (2015).

64. J. D. Lubell, M. H. Brand, Foliar sprays of silver thiosulfate produce male flowers on female hemp plants. HortTechnology 28, 743–747 (2018).

65. A.-M. Faux, X. Draye, M.-C. Flamand, A. Occre, P. Bertin, Identification of QTLs for sex expression in dioecious and monoecious hemp (Cannabis sativa L.). Euphytica 209, 357–376 (2016).

66. S. Han, L. Green, D. J. Schnell, The signal peptide peptidase is required for pollen function in Arabidopsis. Plant Physiol. 149, 1289–1301 (2009).

67. J.-U. Hwang, V. Vernoud, A. Szumlanski, E. Nielsen, Z. Yang, A tip-localized RhoGAP controls cell polarity by globally inhibiting Rho GTPase at the cell apex. Curr. Biol. 18, 1907–1916 (2008).

68. M. Westergaard, The mechanism of sex determination in dioecious flowering plants. Adv. Genet. 9, 217–281 (1958).

69. H. E. Warmke, H. Davidson, Polyploidy investigations. Yearbook of the Carnegie Institution of Washington 1943-44 43, 135–139 (1944).

70. A. Haunold, Cytology, sex expression, and growth of a tetraploid ✕ diploid cross in hop (*Humulus lupulus* L.)^1^. Crop Sci. 11, 868–871 (1971).

71. B. Hyden, J. Zou, D. G. Wilkerson, C. H. Carlson, A. R. Robles, S. P. DiFazio, L. B. Smart, Structural variation of a sex-linked region confers monoecy and implicates GATA15 as a master regulator of sex in Salix purpurea. New Phytol. 238, 2512–2523 (2023).

72. H. C. Wright, C. E. M. Davis, J. Clevenger, W. Korani, KhufuEnv, an auxiliary toolkit for building computational pipelines for plant and animal breeding, Genomics (2025). https://www.biorxiv.org/content/10.1101/2025.03.28.645917v1.

73. D. O. Adams, S. F. Yang, Methionine metabolism in apple tissue: implication of s-adenosylmethionine as an intermediate in the conversion of methionine to ethylene: Implication of S-adenosylmethionine as an intermediate in the conversion of methionine to ethylene. Plant Physiol. 60, 892–896 (1977).

74. D. O. Adams, S. F. Yang, Ethylene biosynthesis: Identification of 1-aminocyclopropane-1-carboxylic acid as an intermediate in the conversion of methionine to ethylene. Proc. Natl. Acad. Sci. U. S. A. 76, 170–174 (1979).

75. T. Boller, R. C. Herner, H. Kende, Assay for and enzymatic formation of an ethylene precursor, 1-aminocyclopropane-1-carboxylic acid. Planta 145, 293–303 (1979).

76. A. J. Hamilton, M. Bouzayen, D. Grierson, Identification of a tomato gene for the ethylene-forming enzyme by expression in yeast. Proc. Natl. Acad. Sci. U. S. A. 88, 7434–7437 (1991).

77. P. Ververidis, P. John, Complete recovery in vitro of ethylene-forming enzyme activity. Phytochemistry 30, 725–727 (1991).

78. A. Boualem, C. Troadec, C. Camps, A. Lemhemdi, H. Morin, M.-A. Sari, R. Fraenkel-Zagouri, I. Kovalski, C. Dogimont, R. Perl-Treves, A. Bendahmane, A cucurbit androecy gene reveals how unisexual flowers develop and dioecy emerges. Science 350, 688–691 (2015).

79. S. Zhang, F.-Q. Tan, C.-H. Chung, F. Slavkovic, R. S. Devani, C. Troadec, F. Marcel, H. Morin, C. Camps, M. V. Gomez Roldan, M. Benhamed, C. Dogimont, A. Boualem, A. Bendahmane, The control of carpel determinacy pathway leads to sex determination in cucurbits. Science 378, 543–549 (2022).

80. C. A. Dowling, J. Shi, J. A. Toth, M. A. Quade, L. B. Smart, P. F. McCabe, S. Schilling, R. Melzer, A FLOWERING LOCUS T ortholog is associated with photoperiod-insensitive flowering in hemp (Cannabis sativa L.). Plant J. 119, 383–403 (2024).

81. R. Vaser, I. Sović, N. Nagarajan, M. Šikić, Fast and accurate de novo genome assembly from long uncorrected reads. Genome Res. 27, 737–746 (2017).

82. A. Astashyn, E. S. Tvedte, D. Sweeney, V. Sapojnikov, N. Bouk, V. Joukov, E. Mozes, P. K. Strope, P. M. Sylla, L. Wagner, S. L. Bidwell, L. C. Brown, K. Clark, E. W. Davis, B. Smith-White, W. Hlavina, K. D. Pruitt, V. A. Schneider, T. D. Murphy, Rapid and sensitive detection of genome contamination at scale with FCS-GX. Genome Biol. 25, 60 (2024).

83. A. M. Bolger, M. Lohse, B. Usadel, Trimmomatic: a flexible trimmer for Illumina sequence data. Bioinformatics 30, 2114–2120 (2014).

84. G. Marcais, C. Kingsford, Jellyfish: A fast k-mer counter. Tutorialis e Manuais, 1–8 (2012).

85. H. Li, B. Handsaker, A. Wysoker, T. Fennell, J. Ruan, N. Homer, G. Marth, G. Abecasis, R. Durbin, 1000 Genome Project Data Processing Subgroup, The Sequence Alignment/Map format and SAMtools. Bioinformatics 25, 2078–2079 (2009).

86. C. Zhou, S. A. McCarthy, R. Durbin, YaHS: yet another Hi-C scaffolding tool. Bioinformatics 39, btac808 (2023).

87. O. Dudchenko, M. S. Shamim, S. S. Batra, N. C. Durand, N. T. Musial, R. Mostofa, M. Pham, B. Glenn St Hilaire, W. Yao, E. Stamenova, M. Hoeger, S. K. Nyquist, V. Korchina, K. Pletch, J. P. Flanagan, A. Tomaszewicz, D. McAloose, C. Pérez Estrada, B. J. Novak, A. D. Omer, E. L. Aiden, The Juicebox Assembly Tools module facilitates de novo assembly of mammalian genomes with chromosome-length scaffolds for under $1000, BioRxiv (2018). 10.1101/254797.

88. J. T. Lovell, A. Sreedasyam, M. E. Schranz, M. Wilson, J. W. Carlson, A. Harkess, D. Emms, D. M. Goodstein, J. Schmutz, GENESPACE tracks regions of interest and gene copy number variation across multiple genomes. Elife 11 (2022).

89. C. J. Grassa, G. D. Weiblen, J. P. Wenger, C. Dabney, S. G. Poplawski, S. Timothy Motley, T. P. Michael, C. J. Schwartz, A new Cannabis genome assembly associates elevated cannabidiol (CBD) with hemp introgressed into marijuana. New Phytol. 230, 1665–1679 (2021).

90. N. Huang, H. Li, compleasm: a faster and more accurate reimplementation of BUSCO. Bioinformatics 39 (2023).

91. L. Gabriel, K. J. Hoff, T. Brůna, M. Borodovsky, M. Stanke, TSEBRA: transcript selector for BRAKER. BMC Bioinformatics 22, 566 (2021).

92. J. M. Flynn, R. Hubley, C. Goubert, J. Rosen, A. G. Clark, C. Feschotte, A. F. Smit, RepeatModeler2 for automated genomic discovery of transposable element families. Proc. Natl. Acad. Sci. U. S. A. 117, 9451–9457 (2020).

93. A. F. A. Smit, R. Hubley, P. Green, RepeatMasker Open-4.0. 2013--2015. [Preprint] (2015).

94. K. Gasic, A. Hernandez, S. S. Korban, RNA extraction from different apple tissues rich in polyphenols and polysaccharides for cDNA library construction. Plant Mol. Biol. Rep. 22, 437–438 (2004).

95. D. Kuznetsov, F. Tegenfeldt, M. Manni, M. Seppey, M. Berkeley, E. V. Kriventseva, E. M. Zdobnov, OrthoDB v11: annotation of orthologs in the widest sampling of organismal diversity. Nucleic Acids Res. 51, D445–D451 (2023).

96. R. van Velzen, R. Holmer, F. Bu, L. Rutten, A. van Zeijl, W. Liu, L. Santuari, Q. Cao, T. Sharma, D. Shen, Y. Roswanjaya, T. A. K. Wardhani, M. S. Kalhor, J. Jansen, J. van den Hoogen, B. Güngör, M. Hartog, J. Hontelez, J. Verver, W.-C. Yang, E. Schijlen, R. Repin, M. Schilthuizen, M. E. Schranz, R. Heidstra, K. Miyata, E. Fedorova, W. Kohlen, T. Bisseling, S. Smit, R. Geurts, Comparative genomics of the nonlegume Parasponia reveals insights into evolution of nitrogen-fixing rhizobium symbioses. Proc. Natl. Acad. Sci. U. S. A. 115, E4700–E4709 (2018).

97. S. Ou, A. Scheben, T. Collins, Y. Qiu, A. S. Seetharam, C. C. Menard, N. Manchanda, J. I. Gent, M. C. Schatz, S. N. Anderson, M. B. Hufford, C. N. Hirsch, Differences in activity and stability drive transposable element variation in tropical and temperate maize. Genome Res. 34, 1140–1153 (2024).

98. M. R. Vollger, P. Kerpedjiev, A. M. Phillippy, E. E. Eichler, StainedGlass: interactive visualization of massive tandem repeat structures with identity heatmaps. Bioinformatics 38, 2049–2051 (2022).

99. G. Benson, Tandem repeats finder: a program to analyze DNA sequences. Nucleic Acids Res. 27, 573–580 (1999).

100. T. L. Madden, The BLAST Sequence Analysis Tool. The NCBI handbook (2013).

101. P. Neumann, A. Navrátilová, A. Koblížková, E. Kejnovský, E. Hřibová, R. Hobza, A. Widmer, J. Doležel, J. Macas, Plant centromeric retrotransposons: a structural and cytogenetic perspective. Mob. DNA 2, 4 (2011).

102. D. M. Emms, S. Kelly, OrthoFinder: solving fundamental biases in whole genome comparisons dramatically improves orthogroup inference accuracy. Genome Biol. 16, 157 (2015).

103. D. M. Emms, S. Kelly, OrthoFinder: phylogenetic orthology inference for comparative genomics. Genome Biol. 20, 238 (2019).

104. K. Katoh, D. M. Standley, MAFFT multiple sequence alignment software version 7: improvements in performance and usability. Mol. Biol. Evol. 30, 772–780 (2013).

105. A. Stamatakis, RAxML version 8: a tool for phylogenetic analysis and post-analysis of large phylogenies. Bioinformatics 30, 1312–1313 (2014).

106. J. Huerta-Cepas, F. Serra, P. Bork, ETE 3: Reconstruction, Analysis, and Visualization of Phylogenomic Data. Mol. Biol. Evol. 33, 1635–1638 (2016).

107. Z. Zhang, J. Li, X.-Q. Zhao, J. Wang, G. K.-S. Wong, J. Yu, KaKs_Calculator: calculating Ka and Ks through model selection and model averaging. Genomics Proteomics Bioinformatics 4, 259–263 (2006).

108. J. K. Lindeløv, mcp: An R Package for Regression With Multiple Change Points (2020). 10.31219/osf.io/fzqxv.

109. T. Pohlert, M. T. Pohlert, Package “pmcmr.” R package version 1 (2018).

110. B. Rosner, Percentage Points for a Generalized ESD Many-Outlier Procedure. Technometrics 25, 165–172 (1983).

111. H. Li, Aligning sequence reads, clone sequences and assembly contigs with BWA-MEM*, arXiv [q-bio.GN]* (2013). http://arxiv.org/abs/1303.3997.

112. H. Li, A statistical framework for SNP calling, mutation discovery, association mapping and population genetical parameter estimation from sequencing data. Bioinformatics 27, 2987–2993 (2011).

113. K. L. Korunes, K. Samuk, pixy: Unbiased estimation of nucleotide diversity and divergence in the presence of missing data. Mol. Ecol. Resour. 21, 1359–1368 (2021).

114. A. Khan, S. B. Carey, A. Serrano, H. Zhang, H. Hargarten, H. Hale, A. Harkess, L. Honaas, A phased, chromosome-scale genome of “Honeycrisp” apple (Malus domestica). GigaByte 2022, gigabyte69 (2022).

115. Z. Xia, W. Fan, D. Liu, Y. Chen, J. Lv, M. Xu, M. Zhang, Z. Ren, X. Chen, X. Wang, L. Li, P. Zhu, C. Liu, Z. Song, C. Huang, X. Wang, S. Wang, A. Zhao, Haplotype-resolved chromosomal-level genome assembly reveals regulatory variations in mulberry fruit anthocyanin content. Hortic. Res. 11, uhae120 (2024).

116. A. Yocca, M. Akinyuwa, N. Bailey, B. Cliver, H. Estes, A. Guillemette, O. Hasannin, J. Hutchison, W. Jenkins, I. Kaur, R. R. Khanna, M. Loftin, L. Lopes, E. Moore-Pollard, O. Olofintila, G. O. Oyebode, J. Patel, P. Thapa, M. Waldinger, J. Zhang, Q. Zhang, L. Goertzen, S. B. Carey, H. Hargarten, J. Mattheis, H. Zhang, T. Jones, L. Boston, J. Grimwood, S. Ficklin, L. Honaas, A. Harkess, A chromosome-scale assembly for “d”Anjou’pear. *G3: Genes, Genomes*, Genetics 14, jkae003 (2024).

117. M. Yang, L. Han, S. Zhang, L. Dai, B. Li, S. Han, J. Zhao, P. Liu, Z. Zhao, M. Liu, Insights into the evolution and spatial chromosome architecture of jujube from an updated gapless genome assembly. Plant Commun. 4, 100662 (2023).

118. L. S. Kubatko, J. H. Degnan, Inconsistency of phylogenetic estimates from concatenated data under coalescence. Syst. Biol. 56, 17–24 (2007).

119. S. V. Edwards, Z. Xi, A. Janke, B. C. Faircloth, J. E. McCormack, T. C. Glenn, B. Zhong, S. Wu, E. M. Lemmon, A. R. Lemmon, A. D. Leaché, L. Liu, C. C. Davis, Implementing and testing the multispecies coalescent model: A valuable paradigm for phylogenomics. Mol. Phylogenet. Evol. 94, 447–462 (2016).

120. M. G. Johnson, E. M. Gardner, Y. Liu, R. Medina, B. Goffinet, A. J. Shaw, N. J. C. Zerega, N. J. Wickett, HybPiper: Extracting coding sequence and introns for phylogenetics from high-throughput sequencing reads using target enrichment. Appl. Plant Sci. 4, 1600016 (2016).

121. A. Bankevich, S. Nurk, D. Antipov, A. A. Gurevich, M. Dvorkin, A. S. Kulikov, V. M. Lesin, S. I. Nikolenko, S. Pham, A. D. Prjibelski, A. V. Pyshkin, A. V. Sirotkin, N. Vyahhi, G. Tesler, M. A. Alekseyev, P. A. Pevzner, SPAdes: a new genome assembly algorithm and its applications to single-cell sequencing. J. Comput. Biol. 19, 455–477 (2012).

122. S. Capella-Gutiérrez, J. M. Silla-Martínez, T. Gabaldón, trimAl: a tool for automated alignment trimming in large-scale phylogenetic analyses. Bioinformatics 25, 1972–1973 (2009).

123. L.-T. Nguyen, H. A. Schmidt, A. von Haeseler, B. Q. Minh, IQ-TREE: a fast and effective stochastic algorithm for estimating maximum-likelihood phylogenies. Mol. Biol. Evol. 32, 268–274 (2015).

124. D. T. Hoang, O. Chernomor, A. von Haeseler, B. Q. Minh, L. S. Vinh, UFBoot2: Improving the ultrafast bootstrap approximation. Mol. Biol. Evol. 35, 518–522 (2018).

125. S. Kalyaanamoorthy, B. Q. Minh, T. K. F. Wong, A. von Haeseler, L. S. Jermiin, ModelFinder: fast model selection for accurate phylogenetic estimates. Nat. Methods 14, 587–589 (2017).

126. C. Zhou, M. Brown, M. Blaxter, The Darwin Tree of Life Project Consortium, S. A. McCarthy, R. Durbin, Oatk: a de novo assembly tool for complex plant organelle genomes, bioRxiv (2024). 10.1101/2024.10.23.619857.

127. N. Dierckxsens, P. Mardulyn, G. Smits, NOVOPlasty: de novo assembly of organelle genomes from whole genome data. Nucleic Acids Res. 45, e18 (2017).

128. R. Bouckaert, T. G. Vaughan, J. Barido-Sottani, S. Duchêne, M. Fourment, A. Gavryushkina, J. Heled, G. Jones, D. Kühnert, N. De Maio, M. Matschiner, F. K. Mendes, N. F. Müller, H. A. Ogilvie, L. du Plessis, A. Popinga, A. Rambaut, D. Rasmussen, I. Siveroni, M. A. Suchard, C.-H. Wu, D. Xie, C. Zhang, T. Stadler, A. J. Drummond, BEAST 2.5: An advanced software platform for Bayesian evolutionary analysis. PLoS Comput. Biol. 15, e1006650 (2019).

129. T. Stadler, Sampling-through-time in birth-death trees. J. Theor. Biol. 267, 396–404 (2010).

130. T. A. Heath, J. P. Huelsenbeck, T. Stadler, The fossilized birth-death process for coherent calibration of divergence-time estimates. Proc. Natl. Acad. Sci. U. S. A. 111, E2957–66 (2014).

131. A. J. Drummond, S. Y. W. Ho, M. J. Phillips, A. Rambaut, Relaxed phylogenetics and dating with confidence. PLoS Biol. 4, e88 (2006).

132. A. Rambaut, A. J. Drummond, D. Xie, G. Baele, M. A. Suchard, Posterior summarization in Bayesian phylogenetics using tracer 1.7. Syst. Biol. 67, 901–904 (2018).

133. S. Magallón, S. Gómez-Acevedo, L. L. Sánchez-Reyes, T. Hernández-Hernández, A metacalibrated time-tree documents the early rise of flowering plant phylogenetic diversity. New Phytol. 207, 437–453 (2015).

134. E, K., H, Monographie Der Früchte Und Samen in Der Kreide von Mitteleuropa (Ústřední ústav geologický v Academii, nakl. Československé akademie věd, 1986).

135. E. M. Friis, P. R. Crane, K. R. Pedersen, Early Flowers and Angiosperm Evolution.,(Cambridge University Press: Cambridge, UK). (2011).

136. M. E. Collinson, The fossil history of the Moraceae, Urticaceae (including Cecropiaceae), and Cannabaceae. Evolution, systematics, and fossil history of the Hamamelidae 2, 319–339 (1989).

137. K. C. Lee, W. Korani, P. C. Bentz, S. Pokhrel, P. Ozias-Akins, A. Harkess, J. Vaughn, J. Clevenger, Long-read Low-Pass sequencing for high-resolution trait mapping, bioRxiv (2025). 10.1101/2025.01.09.632261.

138. N. G. C. Smith, A. Eyre-Walker, Adaptive protein evolution in Drosophila. Nature 415, 1022–1024 (2002).

139. A. McKenna, M. Hanna, E. Banks, A. Sivachenko, K. Cibulskis, A. Kernytsky, K. Garimella, D. Altshuler, S. Gabriel, M. Daly, M. A. DePristo, The Genome Analysis Toolkit: a MapReduce framework for analyzing next-generation DNA sequencing data. Genome Res. 20, 1297–1303 (2010).

140. J. Murga-Moreno, M. Coronado-Zamora, S. Hervas, S. Casillas, A. Barbadilla, iMKT: the integrative McDonald and Kreitman test. Nucleic Acids Res. 47, W283–W288 (2019).

141. J. C. Fay, G. J. Wyckoff, C. I. Wu, Positive and negative selection on the human genome. Genetics 158, 1227–1234 (2001).

142. D. J. Begun, A. K. Holloway, K. Stevens, L. W. Hillier, Y.-P. Poh, M. W. Hahn, P. M. Nista, C. D. Jones, A. D. Kern, C. N. Dewey, L. Pachter, E. Myers, C. H. Langley, Population genomics: whole-genome analysis of polymorphism and divergence in Drosophila simulans. PLoS Biol. 5, e310 (2007).

143. H. Wickham, ggplot2: Elegant Graphics for Data Analysis (Springer, 2016).

144. G. Hickey, J. Monlong, J. Ebler, A. M. Novak, J. M. Eizenga, Y. Gao, Human Pangenome Reference Consortium, T. Marschall, H. Li, B. Paten, Pangenome graph construction from genome alignments with Minigraph-Cactus. Nat. Biotechnol. 42, 663–673 (2024).

145. A. Dobin, C. A. Davis, F. Schlesinger, J. Drenkow, C. Zaleski, S. Jha, P. Batut, M. Chaisson, T. R. Gingeras, STAR: ultrafast universal RNA-seq aligner. Bioinformatics 29, 15–21 (2013).

146. A. R. Quinlan, I. M. Hall, BEDTools: a flexible suite of utilities for comparing genomic features. Bioinformatics 26, 841–842 (2010).

147. M. Pertea, G. M. Pertea, C. M. Antonescu, T.-C. Chang, J. T. Mendell, S. L. Salzberg, StringTie enables improved reconstruction of a transcriptome from RNA-seq reads. Nat. Biotechnol. 33, 290–295 (2015).

148. M. Love, S. Anders, W. Huber, Differential analysis of count data--the DESeq2 package. Genome Biol. 15, 10–1186 (2014).

149. R. Kolde, M. R. Kolde, Package “pheatmap.” R package (2015).

150. R. Poplin, P.-C. Chang, D. Alexander, S. Schwartz, T. Colthurst, A. Ku, D. Newburger, J. Dijamco, N. Nguyen, P. T. Afshar, S. S. Gross, L. Dorfman, C. Y. McLean, M. A. DePristo, A universal SNP and small-indel variant caller using deep neural networks. Nat. Biotechnol. 36, 983–987 (2018).

151. H. Cheng, E. D. Jarvis, O. Fedrigo, K.-P. Koepfli, L. Urban, N. J. Gemmell, H. Li, Haplotype-resolved assembly of diploid genomes without parental data. Nat. Biotechnol. 40, 1332–1335 (2022).

152. G. Marçais, C. Kingsford, A fast, lock-free approach for efficient parallel counting of occurrences of k-mers. Bioinformatics 27, 764–770 (2011).

