## Supplemental Materials for "An X-linked sex determination mechanism in cannabis and hop"

### Supplementary Materials

### Materials and Methods

#### *Genome assemblies*

All sequencing for this study was performed at HudsonAlpha Institute for Biotechnology (Huntsville, AL, USA), unless otherwise noted. To obtain high molecular weight (HMW) DNA for the genome assemblies, we flash froze young leaf tissue in liquid nitrogen then extracted HMW genomic DNA using a Takara NucleoBond HMW kit (San Jose, CA). Libraries were constructed using a SMRTbell Template Prep Kit 3.0, sheared using a Megaruptor (Diagenode), and sized on a Blue Pippin instrument (Sage Science, Beverly, MA, USA). Long-read sequencing was performed using circular consensus sequencing (CCS) mode on a PacBio Sequel II (Pacific Biosciences of California). Sequencing was performed using a 30 hour movie time with two hour pre-extension, and the resulting raw data were processed using the CCS6.2 algorithm. Next, we generated Omni-C data to phase the haplotypes during the assembly process. Libraries were constructed using standard protocols (Cantata Bio, Dovetail Omni-C Kit Catalog #21005). One gram of flash, frozen leaf tissue was ground and after nuclei isolation, the pellet was resuspended in 1x PBS, aliquoted into three tubes and centrifuged at 6000 rpm for five minutes. The supernatant was then discarded and the nuclei pellets flash frozen for storage at -80 degrees. One aliquoted nuclei pellet was then processed using the Dovetail Omni-C protocol V2. Libraries were run on an Illumina NovaSeq 6000 PE150.

For cannabis, HiFi reads were assembled into contigs using hifiasm v0.16.0 (*25*), with the Hi-C integration mode that incorporated Omni-C reads. For hop, we used hifiasm v0.19.5. For 21110M, we also changed the '-s' parameter from 0.55 (default) to 0.2 to better balance the haplotypes out of the assembly. Across all assemblies, we polished our contigs with the HiFi data using Racon v1.5.0 (*81*), removed contigs below 50 Kb using BBTools v38.18-0 (https://sourceforge.net/projects/bbmap/), and removed contigs identified as contamination by FCS-GX v0.4.0 (*82*).

Prior to scaffolding, we manually placed contigs from each of the sex chromosomes into a single haplotype. To accomplish this, we first identified male-specific *k*-mers (Y-mers). We generated Illumina whole-genome sequencing (WGS) for 22 *H. lupulus* individuals (Table S12). DNA was extracted using a CTAB approach and sequencing libraries were constructed using an Illumina TruSeq DNA PCR-free library kit (Catalog #20015962) using standard protocols. Libraries were sequenced on an Illumina NovaSeq 6000 instrument using paired ends and a read length of 150 base pairs. For two isolates (64035M and Bullion) DNA was instead extracted using the BioEcho EchoLUTION Plant DNA 96 Kit (Product No. 010-103-002) and sequencing libraries were constructed using a seqWell purePlex DNA Library Preparation Kit. Sequencing was performed on an Illumina NovaSeq X Plus instrument using paired-end reads and a read length of 150 base pairs at NovoGene (Beijing, China). For hop, we built the k-mer list using 12 of the known sex WGS isolates and the HiFi for two of the genome assembly (*H. lupulus* var.

*lupulus* 21110M and *H. lupulus* var. *lupuloides* MN-1421) to build the Y-mer list. For cannabis we used 14 existing WGS libraries (Table S12). All paired-end Illumina data had adapters removed and were quality filtered using TRIMMOMATIC v0.39 (*83*) with leading and trailing values of three, sliding window of 30, jump of 10, and a minimum remaining read length of 40. We next found all canonical 21-mers in each isolate using Jellyfish v2.3.0 (*84*) and used the bash *comm* command to identify a list of *k*-mers found across all males, but never found in any female, for hop and cannabis separately. We mapped the Y-mers to both haplotypes of each assembly using BWA-MEM, with parameters -k 21 -T 21 -a -c 10 and generated a table of coverage per contig using SAMtools v1.10 *coverage* (*85*). Additional details for manually phasing the sex chromosomes are described in the Supplementary Text.

The sex chromosome-corrected assembly was then scaffolded by first mapping the Omni-C reads using BWA-MEM, with parameters -5SP and samtools v1.10 with parameters view -S -h -b -F 2316. Next, we scaffolded the assembly using Yet Another Hi-C Scaffolding tool (YaHS) v1.1 with default parameters (*86*). We edited contact maps manually with Juicebox Assembly Tools (JBAT) v2.15 (*87*) to produce the expected 10 chromosomes per haplotype. Scaffolds containing assembled telomeres were checked for correct orientation to the terminal ends by searching for the $(TTTAGGG)_n$ repeat using GENESPACE v1.3.1 (*88*). The cannabis assemblies were reordered and oriented to the cs10 reference (*89*) and hop were reordered and oriented to Cascade (*17*). Final plots showing the number of contigs and telomeric regions (density of the window for the telomeric repeat set to 0.75) were generated using GENESPACE. We assessed the quality of the genome assemblies using compleasm v0.2.6 (*90*) using the Eudicots database and Merqury v1.3 (*26*).

***Gene annotations***

To generate gene model annotations, for cannabis, we annotated repeats and genes following previously described methods (*58*). In brief, we used a repeat library built from existing assemblies and repeats were masked with RepeatMasker v4.1.2. We annotated gene models using Braker v2.1.6 and TSEBRA (*91*), using RNAseq and Full-length Oxford Nanopore cDNA (Table S12).

For the hop assemblies, we annotated repeats by building a *de novo* library for each haplotype separately using RepeatModeler v2.0.5 (*92*), invoking the -LTRStruct parameter. We then masked repeats using RepeatMasker v4.1.5 (*93*). We annotated gene models by using BRAKER3 v3.0.6 (*35*), using both proteins and RNAseq data as extrinsic evidence. To generate RNAseq data, we flash froze leaf tissue and inflorescences for *H. lupulus* var. *lupulus* 21110M and Comet "male", respectively. We extracted RNA using a CTAB approach (*94*). Sequencing libraries were constructed using an Illumina TruSeq Stranded mRNA Library Prep Kit (Catalog #20020595) using standard protocols. Libraries were sequenced on an Illumina NovaSeq 6000 instrument using paired ends and a read length of 150 bp. We additionally used existing RNAseq data (Table S12). For the protein support, we used the OrthoDB v11 (*95*) for Viridiplantae, combined with proteins from genome references for *Parasponia (96)*, *Trema (96)*, Cascade, and cs10. We assessed the gene annotations using compleasm v0.2.6 (*90*) using the Eudicots database.

***Repeat analyses***

To examine TEs, we additionally annotated repeats using EDTA v2.0.0 (*43*) in the sensitive mode for each haplotype separately. To compare repeat landscapes between cannabis and hop, we generated a panEDTA library using EDTA v2.2.0 (*97*). We next used this pan-library to identify the repeats on the sex chromosomes using RepeatMasker, and Kimura substitution values were calculated using the script createRepeatLandscape.pl.

To identify putative centromere locations, we used StainedGlass v0.5(*98*) to visualize the massive tandem repeat arrays. To identify tandem repeats, we used Tandem Repeats Finder v4.09.1 (*99*) (parameters 2 7 7 80 10 50 500 -f -d -m -h) and to identify CRM elements we used BLAST v2.15.0 (*100*) using previously identified elements (*101*). To plot the landscape of repeats and genes in the context of the putative centromere locations, we use sliding windows of 100 Kb and 200 Kb for the Y and X, respectively.

***Synteny analyses***

For cannabis and hop, visualization and analysis of local synteny was conducted with DEEPSPACE (https://github.com/jtlovell/DEEPSPACE), which aligns genomic windows between assemblies and uses GENESPACE machinery to infer syntenic blocks. Source code for the riparian plots was modified where needed to visually compare two genomes with such different total sizes. To examine syntenic relationships between Cannabaceae and Moraceae, we use GENESPACE using default parameters.

***Gene tree analyses***

To compare the boundary of the MSY with the PAR when using Y-mer mapping, and examine the timing of gene capture into the MSY, we built 1,096 gene trees that contained a Y-linked gene model. We first ran OrthoFinder v2.5.2 (*102*, *103*) in ultra-sensitive mode, with the cannabis and hop proteins and 42 additional angiosperms (Table S13). We aligned the genes using MAFFT v7.471 (*104*) with the parameter maxiterate set to 1000 and using genafpair. We built gene trees using RAxML v8.2.12 (*105*) with 100 bootstrap replicates and invoking the model PROTGAMMAWAG. We visually examined the topology to assess whether sex-linkage of the gene occurred prior to, or after, the divergence of the genera. To examine phylogenetic evidence of monoecy evolving from a Y-linked gene translocating to the X, we used the ETE3 v3.1.1 (*106*) *check_monophyly* function. This OrthoFinder run was also used to examine gene homology and due to copy number variation, gene numbers given in the main text are from the cannabis ‘Otto II’ reference.

***Ks-based analyses and age estimation for the sex chromosomes***

To identify one-to-one orthologs on the sex chromosomes (gametologs), we ran OrthoFinder v2.5.2 within a reference's haplotypes. We calculated synonymous (Ks) changes in codons using Ka/Ks Calculator v2.0 (*107*). To test for evidence of multiple capture events into the MSY (i.e., evolutionary strata), we used the R package mcp v0.3.4 (*108*) on Ks values. In cannabis, we excluded one outlier (Ks of 3.7) identified using PMCMRplus v1.9.12 (*109*), which runs Rosner's generalized extreme studentized deviate many-outlier test (*110*). For mcp, we used the model, 'y ~ 1, ~ 1' to identify the change point between two plateaus, and we used 10,000 iterations, 3 chains, and a burn-in of 10,000 (i.e., 'adapt'). We also tested the model 'y ~ 1, ~ 1, ~1' and 'y ~ 1, ~ 1, ~1' to search for a third and fourth plateau, respectively.

To estimate the age of the sex chromosomes, we used the following the formula: time (T)= $(Ks/\boldsymbol{\pi}-1)x(2N_e t_g)$. We used values from hop to estimate the age. For Ks, we used both the max Ks (0.4), as well as the average of the top 10% of Ks values (0.33). We used a $\mu$ of $1x10^{-9}$ and a generation time ($t_g$) of two. For $\boldsymbol{\pi}$, we used 0.016, which we calculated by first using BWA-MEM v0.7.17 (*111*) to map the WGS data to the *H. lupulus* var. *lupulus* HAP1 reference, where we removed the Y chromosome, but included the X chromosome from HAP2. We used bcftools v1.9 *mpileup* and *call (112)* functions to call variants, where we applied the -G parameter for *call*. We filtered the vcf file using 'QUAL>20 & DP>5 & MQ>30', minor allele frequency of 0.05, and dropped sites with > 25% missing data. We identified SNPs using all 22 *H. lupulus* genotypes generated here, and one existing *H. scandens* genotype; however, we restricted the estimation of $\boldsymbol{\pi}$ to wild-collected female hop genotypes (N=7). Last, $\boldsymbol{\pi}$ was estimated using pixy v2.0.0.beta8 (*113*) with 100,000-bp windows and a 10,000-bp jump, and the average taken across the autosomes only.

***Species tree inference***

We used OrthoFinder v2.5.4 (*102*, *103*) to identify conserved, single-copy genes for species tree inference with ASTRAL v5.7.8 (*36*), which employs a coalescent-based summary method that is statistically consistent under the multi-species coalescent model. To identify orthologs conserved as single copy across the Rosales order, we used gene annotations from the following Rosales taxa: *H. lupulus* var. *lupuloides*, *H. lupulus* var. *lupulus*, *H. lupulus* var. *neomexicanus*, *C. sativa*, *Ficus hispida (31)*, *Malus domestica (114)*, *Morus alba (115)*, *Pyrus commune (116)*, and *Ziziphus jujuba (117)*. All single-copy genes conserved across those samples were used for gene and species tree inference, after removing all orthologs of sex chromosome linked genes (found in any species) since any of those may exhibit unique evolutionary histories in different lineages. Coalescent-based, rather than concatenation, methods were used to analyze nuclear genes because concatenation implicitly assumes an absence of recombination and ILS, or incomplete lineage sorting (i.e., a single history is shared amongst all genes), and can result in statistical inconsistencies in multi-locus datasets with high levels of gene tree discordance (*118*, *119*); whereas coalescent approaches assume free recombination between genes, accounting for gene tree–species tree discordance due to ILS.

Along with our *Humulus* dataset, publicly available Illumina short reads were downloaded from NCBI SRA for additional lineages spanning the Rosales (Table S14), to produce gene assemblies for phylogenomic analysis. We employed HybPiper v2.1.6 (*120*) to i) map reads to target nucleotide sequences with BWA-MEM, ii) assemble mapped reads into contiguous sequences with SPAdes v3.15.2 (*121*), iii) extend gene assemblies into flanking intronic regions, and iv) test for paralogs with default HybPiper parameters. The target sequence file used for read mapping with HybPiper was manually curated with nucleotide coding sequences for 775 one-to-one orthologs from genome annotations for: 1) *H. lupulus* var. *neomexicanus*, 2) *M. domestica*, 3) *C. sativa*, 4) *M. alba*, and 5) *Z. jujuba*.

Multiple sequence alignments for each target gene were produced using MAFFT v7.520 with the *--auto* option, which automatically selects the best alignment strategy based on the data (*104*). Poorly-aligned sequences were trimmed using trimAl v1.4.1 with the flag *-automated1* (*122*). Gene tree inference was performed using maximum likelihood (ML) with IQ-TREE v1.6.2 (*123*), 1000 ultrafast bootstraps (BS) (*124*), and model selection based on Bayesian information criterion with ModelFinder (*125*). Prior to the ASTRAL analysis, gene tree branches with <10% BS support were collapsed into polytomies to improve the accuracy of species tree inference (*36*). The resulting species tree was rooted with Rosaceae taxa (i.e., *P. communis*, *M. domestica*).

***Plastome divergence time estimations***

Whole (circularized) plastome assemblies were used to estimate relationships and divergence times among the same, or similar, Rosales taxa included in the species tree analysis (Table S14). To assemble chloroplast genomes, we used oatk v0.2 (*126*) when HiFi data was available. For all others where we used Novoplasty v4.3.5 (*127*). Limited taxon sampling was used to optimize time calibrations across the Rosales tree, and obtain robust divergence time estimates for ingroup branches (i.e., Cannabaceae and *Humulus*). Whole plastomes were aligned using MAFFT v7.520 with the *--auto* option and used as input for ML tree inference with IQ-TREE v1.6.2 (as performed for individual nuclear gene trees) and bayesian divergence time estimation with BEAST v2.7.7 (*128*). BEAST was executed using a GTR+G+I substitution and site heterogeneity model with four rate categories; a fossilized birth-death (FBD) tree prior to model speciation and extinction (*129*, *130*); and optimized relaxed clock model with a lognormal distribution (*131*). The plastome tree topology was constrained according to the ML tree from IQ-TREE and rooted with Rosaceae taxa. Two independent BEAST runs, totalling 48,858,000 MCMC generations, were combined to obtain >200 effective sample size (ESS) for all priors, with MCMC sampling every 1000 generations and 10% burn-in. ESS was assessed with Tracer v1.7.2 (*132*). To calibrate the plastome tree to time (i.e., million years ago=Ma) and estimate the tree root height, we estimated the FBD origin under a normal distribution with a mean of 107 Ma and a liberal standard deviation of 4.0. Those parameters virtually constrained the total tree height to approximately 100-114 Ma, which is in accordance with crown group age estimates for Rosales in previous studies (*28*, *133*). Under the FBD model, two fossil calibrations were applied to further calibrate the plastome tree to time: the first applied a 66 Ma minimum age for the

Cannabaceae stem group (*28*, *134*, *135*) and the second applied a 33.9 Ma minimum age for the stem group of *Humulus* (*28*, *136*).

***Tests for selection***

To examine protein evolution on the X chromosomes, and test for evidence of faster-X evolution, we first identified OrthoGroups using OrthoFinder v2.5.2 with HAP2 of *H. lupulus* var. *lupulus*, *H. lupulus* var. *neomexicanus*, and *C. sativa* 'Otto II', as well as *Morus notibilis (32)*, and *Trema orientale (96)*. We found 10,056 single-copy OrthoGroups where all five genomes were represented. To calculate the ratio of nonsynonymous to synonymous changes in proteins (dN/dS), we used codeml in PAML v4.10.7 (*44*). We assessed significance between autosomal, PAR, and HXR genes, where we excluded dN/dS values greater than 3, and ran a Wilcoxon rank sum test with a Benjamini Hochberg correction for multiple tests.

To calculate Fst and identify outlier regions between chemotypes I-II and III-IV, and between female and monoecious genotypes in cannabis, we generated PacBio HiFi long-read sequencing for 12 additional monoecious genotypes. For two samples, (NWG_2624-03-H and NWG_3584-03-H), the methods for generating HiFi were the same as described above. For the 10 remaining genotypes, seeds were sown in greenhouse conditions before being transplanted to an outdoor (in-ground) plot in Geneva, New York (USA). Young leaf tissue was collected and flash frozen on liquid nitrogen. We extracted HMW DNA, prepared and pooled PacBio HiFi libraries, and sequenced using the PacBio Revio platform as described by (*137*). Plant sex phenotypes were recorded for each individual on a weekly basis throughout the growing season. We additionally used 51 existing HiFi datasets (Table S12) (*58*). We aligned the reads to the 'Otto II' HAP2 reference using pbmm2 v.1.17.0 (https://github.com/PacificBiosciences/pbmm2). To generate an all-sites VCF, we called SNPs using bcftools *mpileup* and *call*. We filtered the vcf file using 'QUAL>20 & DP>3 & MQ>30', a minor allele frequency of 0.05, and dropped sites with > 25% missing data. We used pixy v2.0.0.beta8 (*113*) using a 100,000-bp window with a 10,000-bp jump. We defined outliers as three standard deviations above the mean, making the outliers above 43.8 for female versus monoecious comparisons and 20.6 for between chemotypes.

To estimate adaptive substitutions, or alpha ($\alpha$) (*138*), in hop we used the McDonald Kreitman Test (MKT). We first identified variants as described above. Next, we used the GATK v4.5 (*139*) FastaAlternateReferenceMaker tool to convert the VCF for each genotype to fasta format and used gffread v0.12.7 (https://github.com/gpertea/gffread) to output coding sequences. We used the R package iMKT v0.1.1 (*140*), where we applied a Fay, Wyckoff, and Wu correction (*141*) of 0.15. Because the MKT can be underpowered when there are low counts in the 2-by-2 contingency table, we only compared results for which the row and column totals were six or greater (*142*). After calculating the MKT, we used the Fisher exact test to test for significance on each gene and applied a false discovery rate adjustment to the p-values.

***Backcross generation and phenotyping***

To generate the $BC_1$ population, a cross was made between New West Genetics' breeding lines using NWG 7619 as the female and NWG 7384 as the sire. NWG 7619 is an early flowering dioecious variety and NWG 8232 is a later flowering monoecious variety. Crossing was performed in a greenhouse in Fort Collins, CO where plants were grown in 3.5" pots using Pro-Mix HP potting soil with supplemental lighting set for 16-hour daylength. A monoecious NWG 7384 was paired with a NWG 7619 female under a Midco (St. Louis, MO) pollination bag. F1 seed was then planted outdoors in a small plot in Winsor, CO where XY males were rogued so that only NWG 8232 pollen was available to sire $BC_1$ seed on F1 females. Approximately 20 grams of $BC_1$ seed was harvested from two F1 plants. $BC_1$ seed was planted in a greenhouse under conditions similar to those already described and used for the phenotyping the degree of monoecy using the following 0 to 4 scale: 0 = no staminate flowers, 1 = more pistillate than staminate, 2 = roughly equal pistillate and staminate flowers, 3 = more staminate than pistillate, 4 = only staminate flowers. Floral phenotypes were scored as $BC_1$ plants flowered over the course of an approximately 4 week period.

To ensure that no Y chromosomes were present in the $BC_1$ data, which may impact the results, we tested for the presence of the Y chromosome. To accomplish this, we used the *bbduk* function of BBmap v38.86 (Bushnell, sourceforge.net/projects/bbmap/) to identify reads that contained perfect matches to the list of cannabis Y-mers. We mapped these reads to the Y-containing HAP1 of the 'Otto II' reference, using BWA v0.7.17 (*111*) and we used Samtools v.1.10 (*85*) to calculate coverage of the non-recombining region. We plotted these coverage values using ggplot2 (*143*) and found no evidence of a Y chromosome (Fig. S11).

KhufuPAN (https://github.com/w246 korani/KhufuPAN) was used to produce a genome graph and analyze low coverage whole-genome sequencing. A graph was constructed using Minigraph-cactus (*144*) with each haplotype individually in the graph of 'Carmagnola', 'Futura 75', 'Otto II', and 'Uso 31'. The reference genome used was HAP2 of 'OttoII' with the X. The graph consists of 8,303,397 SNPs and 2,610,350 SVs. A total of 950 lines (average depth 1.01X) were mapped to the pangraph and variants were genotyped. After filtering for minimum depth of 2, missing data across lines less than 75%, and minor allele frequency of 0.01, 906,075 variants were retained in the panmap. Filtering for lowly sequenced lines resulted in retaining 863 lines for the final analysis.

Identification of significant allele frequency differences between male and female classes was done using KhufuEnv (*72*). The genotypes from the female (N=367) and monoecious (N=520) samples were extracted, group-specific allele frequency was calculated, and the difference in allele frequency was calculated. Significant sites with allele frequency difference of greater than 0.4 were identified.

***Gene expression analyses***

To examine gene expression in cannabis, we used existing RNAseq data for leaf, stem, and apical meristem tissues (N=72; Table S12) (*54*, *80*). We first trimmed the data as described above, and aligned the data using STAR v.2.7.9a (*145*) to the 'Otto II' HAP2 reference, including the Y. To inhibit multi-mapping issues with the PAR being represented twice on the XY chromosomes, we hard-masked the Y PAR using bedtools v2.31.0 *maskfasta* (*146*). We assembled transcripts using StringTie v2.1.7 (*147*) and ran differential gene expression using DESeq2 v1.46 (*148*). To generate the heatmap of expression, we used 28 additional RNAseq libraries across tissue types (N=100 total). We generated the heatmap using pheatmap v1.0.12 (*149*) in R v4.4.2.

***Additional variant calling***
In addition to identifying SNPs with bcftools *mpileup* in the cannabis HiFi data, we also called SNPs and small INDELs using DeepVariant v1.8.0 (*150*) and larger SVs using Sawfish v0.12.10 (https://github.com/PacificBiosciences/sawfish).

### Supplementary Text

***Sex chromosome manual curation***
Sex chromosomes can be particularly difficult to assemble in a *de novo* genome reference. This difficulty is partially due to computational challenges of phasing the sequences: without parental information (e.g., trio-bins) the X/Y chromosomes can end up partially represented across both haplotypes. This of course also affects autosomes (*1*), but the highly divergent and structurally variable nature of some sex chromosomes (*2*) can make scaffolding extremely difficult. Without trio-binning there are two extremes (and many possible variants in between) for the sex chromosomes. If the male-specific region of the Y (MSY) is contained in a single contig, one can move straight on to scaffolding the assemblies. The other extreme is when the MSY is split between haplotypes (e.g., X-linked and Y-linked contigs are found in both haplotype 1 and 2). In these cases, the X and Y-linked contigs will need to be identified and manually moved to a single haplotype each. As such, we have developed an approach to phase the sex chromosomes that relies on the generation of sex-linked *k*-mers (*3*) to identify X- versus Y-linked contigs.

In the *Cannabis* and *Humulus* assemblies, to determine first in which haplotype were the Y-linked contigs, we used whole-genome Illumina data of known sex isolates (and PacBio HiFi for the *H. lupulus* var. *lupulus* 21110M and *H. lupulus* var. *lupuloides* MN-1421 isolates only). All paired-end Illumina data had adapters removed and were quality filtered using TRIMMOMATIC v0.39 (*4*) with leading and trailing values of 3, sliding window of 30, jump of 10, and a minimum remaining read length of 40. We next found all canonical 21-mers in each isolate using Jellyfish v2.3.0 (*5*) and used the bash *comm* command to find all *k*-mers shared in all male isolates and not found in any female isolate (Y-mers). A similar approach can be taken using meryl (*6*), which can generate the k-mer *counts*, and has *intersect* and *difference* commands for finding male-specific k-mers. We mapped the Y-mers to both haplotype assemblies using BWA-MEM v0.7.17 (*7*), with parameters '-k 21' '-T 21' '-a' '-c 10' and generated a table of coverage per contig using SAMtools v1.10 *coverage* (*8*). Y-linked contigs have high Y-mer coverage (though in our experience coverage ~1 or greater can suggest

Y-linkage; even lower values are possible) and/or have a high proportion of bases covered. The exception is for the pseudoautosomal region (PAR); an entirely PAR-derived contig will not show a density of Y-mers, but if a portion is contiguously assembled with a piece of the MSY, some Y-mer coverage is expected. The coverage and basepair cutoffs for species outside of *Cannabis* and *Humulus* may vary based on the complexity of the MSY and factors that could affect the Y-mer list (some discussion here (*3*)). We found that for the *H. lupulus* var. *lupuloides* reference, the Y chromosome assembled in a single contig, so could be scaffolded directly (Table S2). For the remaining *Cannabis* and *Humulus* isolates, we found evidence that the Y chromosome was assembled into several contigs, and in most cases split between the haplotypes (Table S2).

To identify X-linked contigs, and additionally validate the Y-linked contigs using a second approach, we used a "joint-scaffolding" approach, where we concatenated the two haplotypes from the assembly together and used Yet Another Hi-C Scaffolding tool (YaHS) (*9*) to scaffold it. We note that this joint-scaffolded assembly was not used for downstream analyses, only for sex-linked contig identification. To accomplish the scaffolding, we mapped the Omni-C reads using BWA-MEM v0.7.17 with parameters '-5SP' and samtools v1.10 with parameters view '-S' '-h' '-b' '-F 2316'. Next, we scaffolded the assembly using YaHS v.1.1 with default parameters. We visually examined the contact maps with Juicebox Assembly Tools (JBAT) v2.15 (*10*) and manually curated the Y and X chromosomes. For the Y-linked contigs, we first searched for all the contigs identified with Y-mers (Table S2) and visually examined the contacts. We then searched for contacts that were not identified with our Y-mers, which facilitated the identification of the PAR contigs for the Y. Using the PAR, we search for "offdiagnonal" contacts that represent the PAR homology to identify the X chromosome (Fig. S10) and again leveraged contacts with the X to further identify X-linked contigs. For *Cannabis* and *Humulus*, a Y-linked contig was moved into HAP1 if it showed Y-mer coverage >0.9 (Table S2) and scaffolded contiguously (including with manual curation). X-linked contigs were moved into HAP2 if they scaffolded contiguously (including with manual curation) (Fig. S10). It is worth also noting that here we have focused on XY sex determination mechanisms, but the logic applies equally to ZW systems (*3*).

***Divergence time estimation***

Divergence time estimates from this study, based on whole plastomes, were generally in agreement with similar previous analyses (Table S15) that employed few nuclear and/or organellar markers (Magallón et al., 2015; Zhang et al., 2018; Jin et al., 2019). Focussing on clade age estimates relevant to this study, our estimated 95% highest posterior density (HPD) intervals were at least partially overlapping with most 95% HPD intervals from those previous studies, aside from two nodes in Jin et al. (2019) (Table S.divTime). The differences observed amongst clade age estimates from this study and the others presented in Table S.divTime are likely due to differences in 1) data analyzed (i.e., whole plastome assemblies in this study vs. few genetic markers in the others); 2) priors/models applied for age estimations; and 3) species samples (e.g., whereas Jin et al. [2019] used Urticaceae and Moraceae as outgroups, in this study, and the other two, Rosaceae or more distant relatives were used for such).

### Supplementary Figures

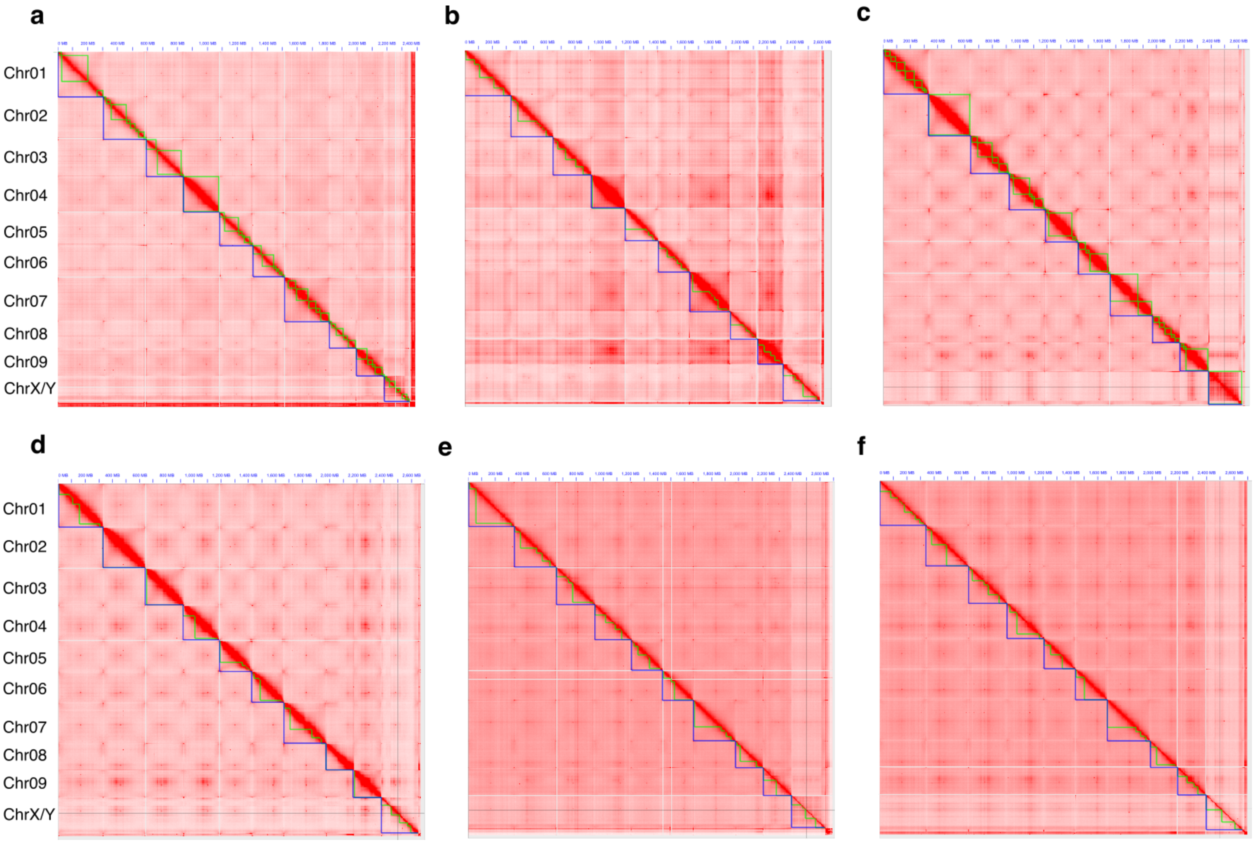


**Fig. S1. Hi-C contact maps for the phased *Humulus* haplotypes.** The Hi-C contact maps show the expected 10 chromosomes for both haplotypes for **a**) *Humulus lupulus* var. *lupulus* HAP1, **b**) *Humulus lupulus* var. *lupulus* HAP2, **c**) *Humulus lupulus* var. *lupuloides* HAP1, **d**) *Humulus lupulus* var. *lupuloides* HAP2, **e**) *Humulus lupulus* var. *neomexicanus* HAP1, **f**) *Humulus lupulus* var. *neomexicanus* HAP2.

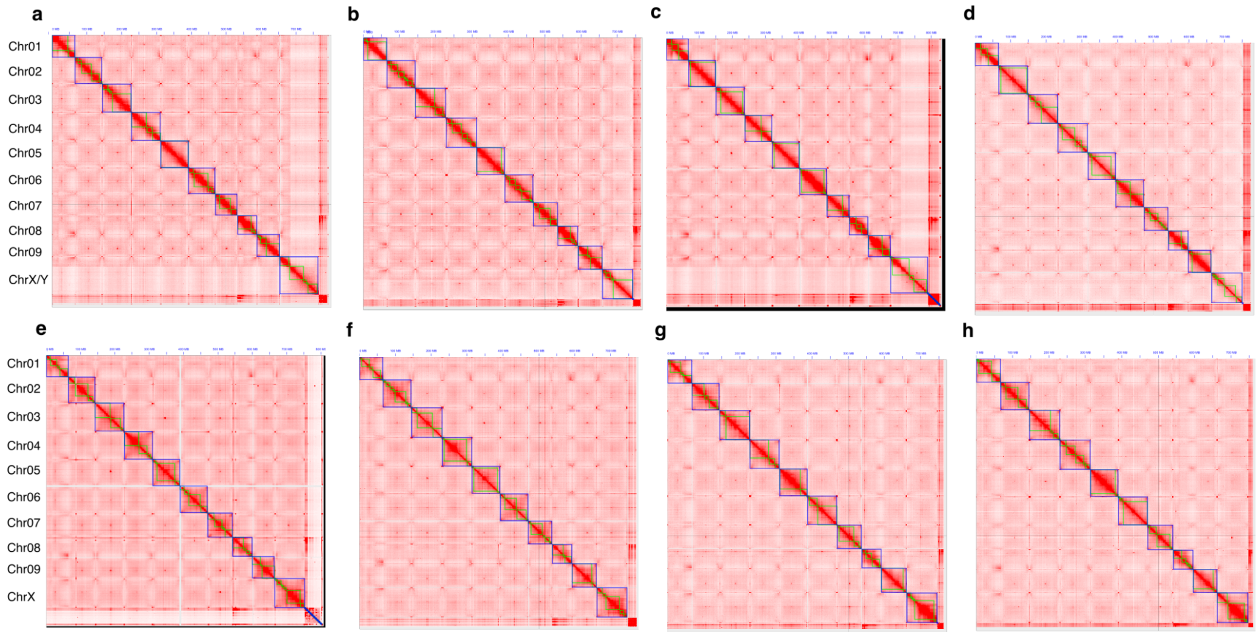


**Fig. S2. Hi-C contact maps for the phased *Cannabis* haplotypes.** The Hi-C contact maps show the expected 10 chromosomes for both haplotypes for **a**) Cannabis sativa cv. Carmagnola HAP1, **b**) Cannabis sativa cv. Carmagnola HAP2, **c**) Cannabis sativa cv. Otto II HAP1, **d**) Cannabis sativa cv. Otto II HAP2, **e**) Cannabis sativa cv. Futura 75 HAP1, **f**) Cannabis sativa cv. Futura 75 HAP2, **g**) Cannabis sativa cv. Uso 31 HAP1, **h**) Cannabis sativa cv. Uso 31 HAP2.

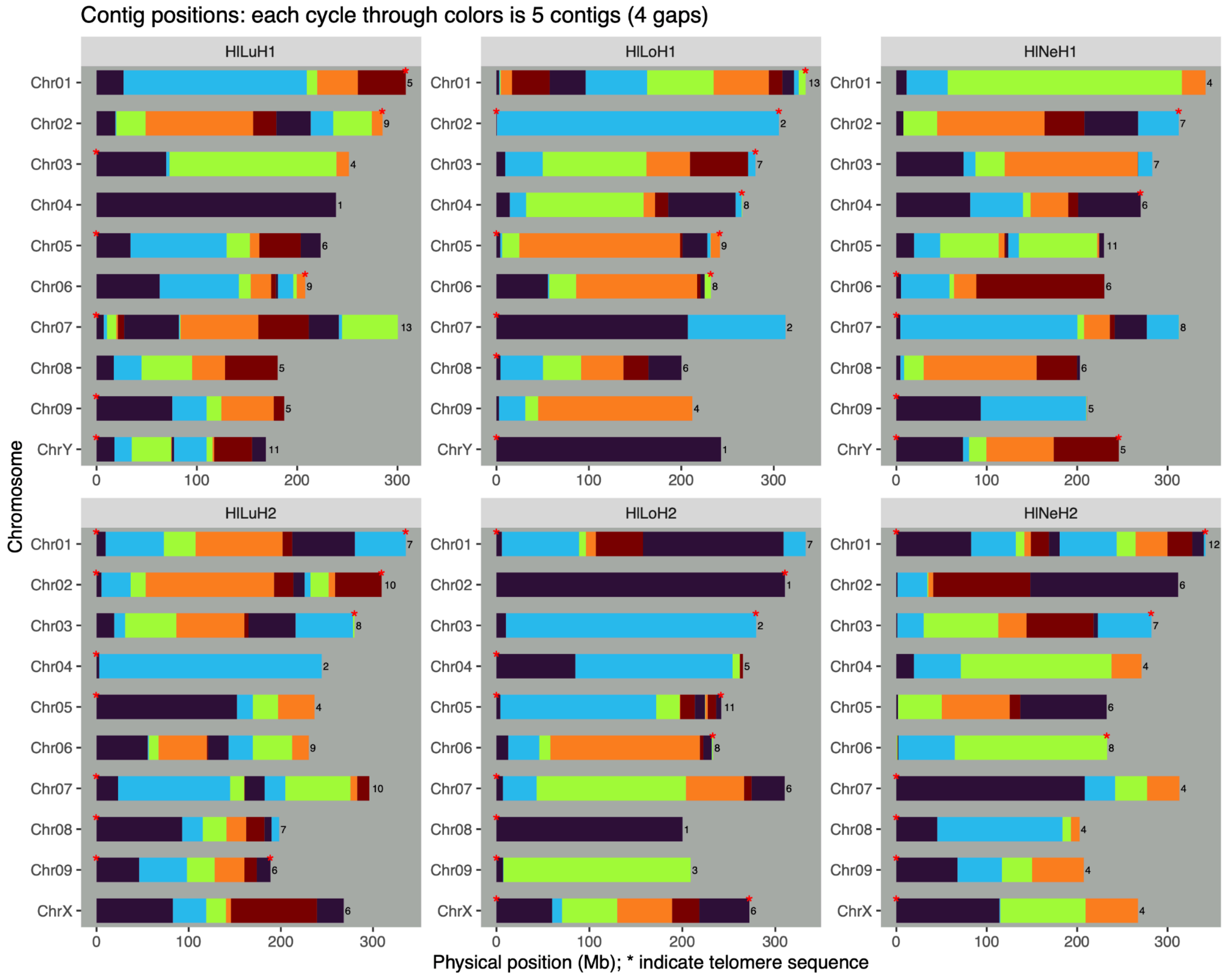


**Fig. S3. Contig map and telomeric sequence location for the *Humulus* genome assemblies.** The contigs in each genome are shown as a continuous block of a single color, cycling through a discrete sequence of five colors. The number of contigs for each chromosome are shown to the right of the final contig. Red asterisks indicate telomere sequences.

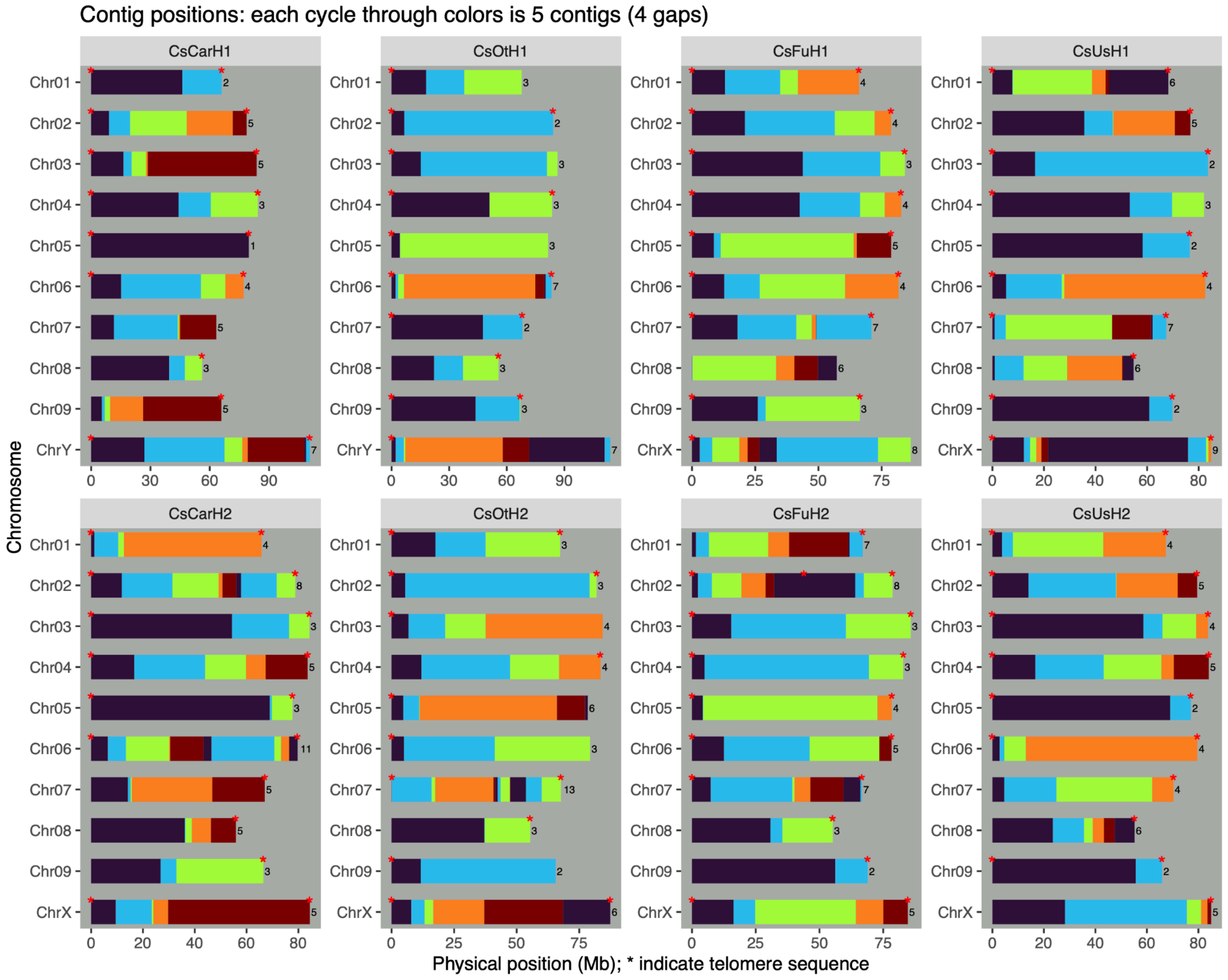


**Fig. S4. Contig map and telomeric sequence location for the *Cannabis* genome assemblies.** The contigs in each genome are shown as a continuous block of a single color, cycling through a discrete sequence of five colors. The number of contigs for each chromosome are shown to the right of the final contig. Red asterisks indicate telomere sequences.

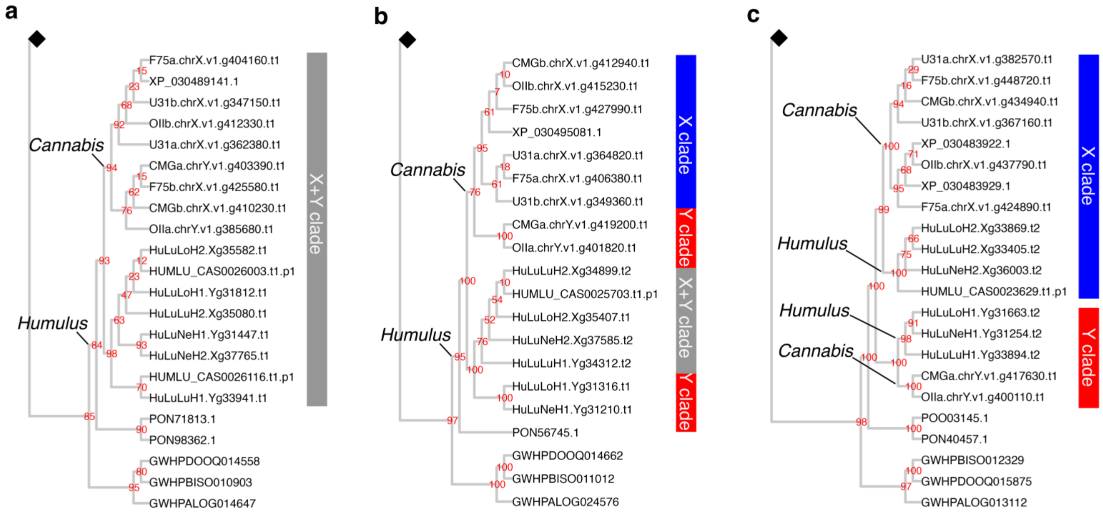


**Fig. S5. Example gene trees with different topologies on the sex chromosomes**. **a**) an example gene tree showing a gene in the pseudoautosomal region, where the topology reflects the species tree and X- and Y-linked genes within the same clade. **b**) an example gene tree showing independent captures of genes in the sex-determination region, as evidenced by the Y-linked genes with *Cannabis* and *Humulus* forming their own clades. **c**) an example gene tree showing sex-linkage prior to the divergence of the genera. This is evident by the X- and Y-linked copies forming a clade, rather than following the species topology in panel a. The numbers in red are bootstrap supports and the black diamond represents where the remaining portion of the tree was collapsed.

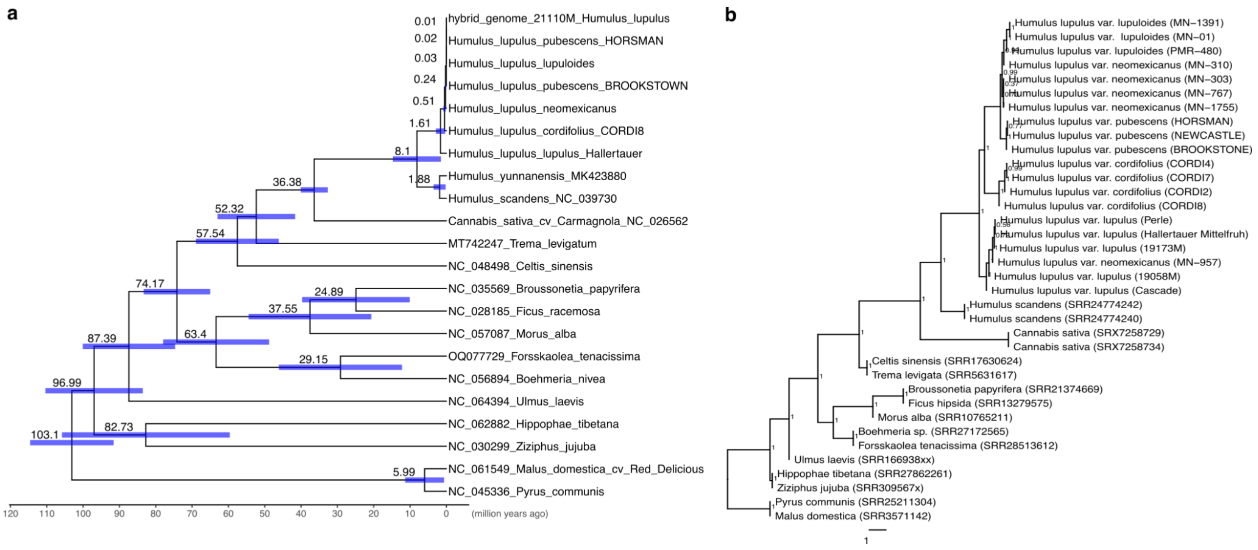


**Fig. S6. Divergence time tree (a) estimated with BEAST (v.2.7.7), based on whole plastome assemblies, and species tree (b) estimated using a coalescent-based summary approach with ASTRAL (v.5.7.8), based on 775 nuclear genes.** Branches and time scale in the BEAST tree are divergence times (million years ago), node labels representing the mean, and blue confidence intervals are 95% highest posterior densities (HPDs). Node labels in the ASTRAL tree are local posterior probabilities and branch lengths correspond to coalescent time units.

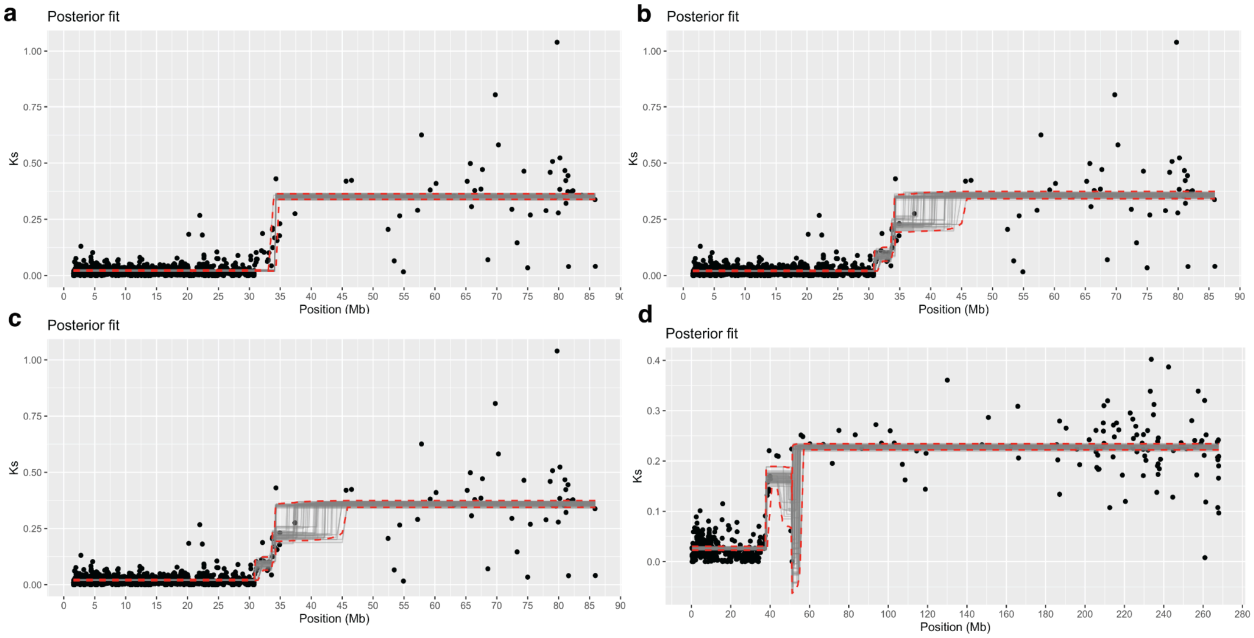


**Fig. S7. Change point analyses to identify the location of strata boundaries. a**) Change point when using Ks of one-to-one XY orthologs in cannabis. The gray lines represent posterior draws, the red dashed lines are the 2.5% and 97.5% quantiles, and the blue line is the posterior density of the change point. In this panel, we tested for a single change point (two plateaus). **b**) testing the change point when the model accounts for two changes (three plateaus). There is evidence for a second change point, within the region of independent captures into the non-recombining region. **c)** testing the change point when the model accounts for three changes (four plateaus) does not support a third change point, and none within the region shared with hops. **c)** testing the change point using hop (allowing three changes) shows the same patterns as in cannabis.

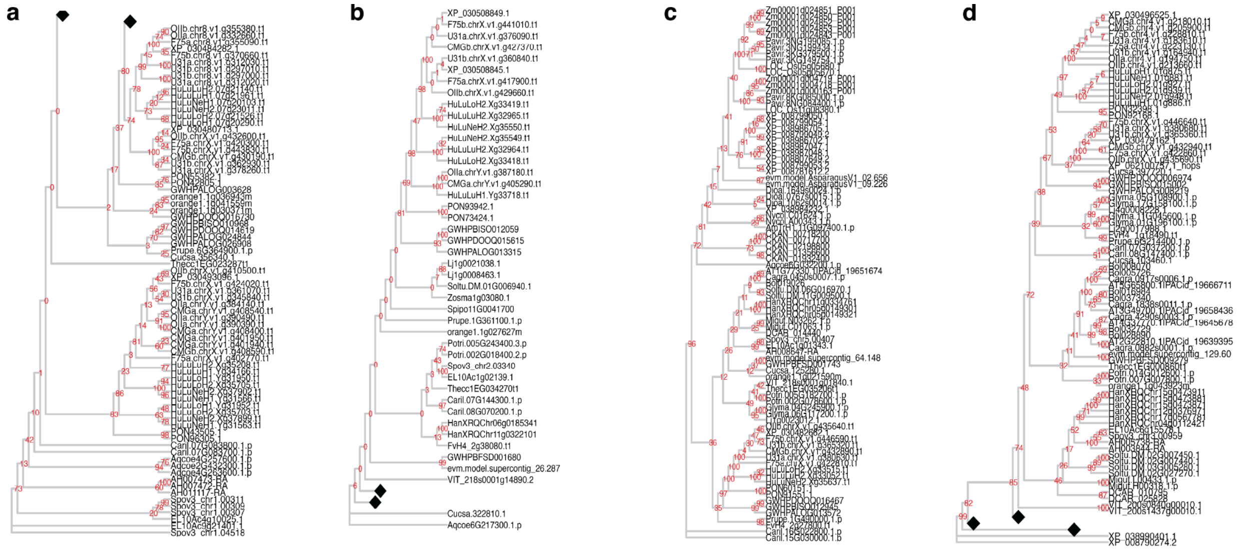


**Fig. S8. Gene trees for select flowering and ethylene genes. a**) Flowering locus T, **b**) Flowering locus D, **c**) Aminocyclopropane-1-carboxylate synthase (ACC) oxidase (ACO); **d**) ACC synthase (ACS). The numbers in red are bootstrap supports and the black diamond represents where the remaining portion of the tree was collapsed.

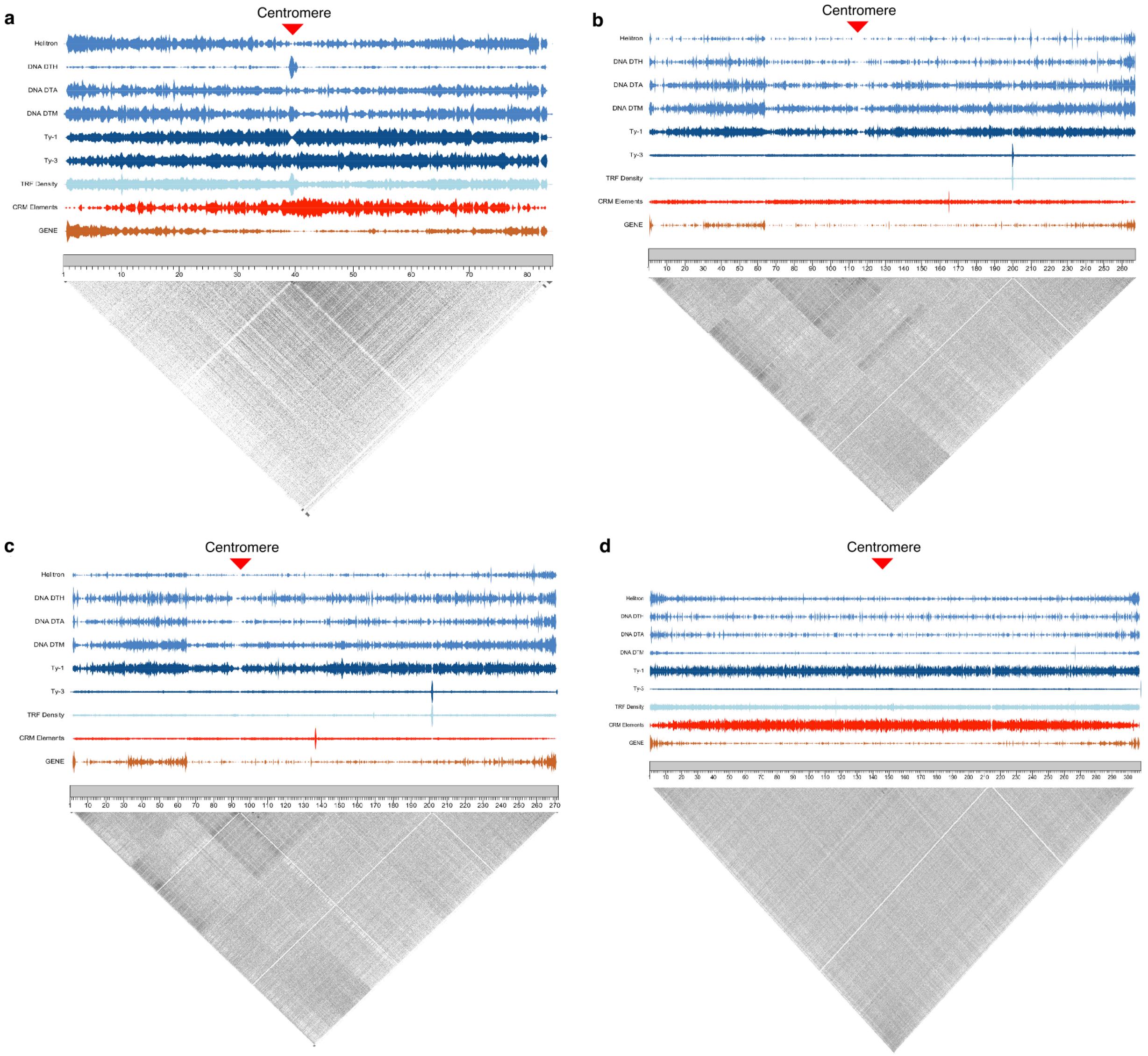


**Fig. S9. Putative centromere locations. a**) putative centromere location for the *Cannabis sativa* cv. Carmagnola X, (**b**) *Humulus lupulus* var. *neomexicanus* X (**c**) *Humulus lupulus* var. *lupuloides* X (**d**) *Humulus lupulus* var. *lupulus* Chr01 and (**b**) X chromosomes showing their different proximities from the pseudoautosomal regions (PAR). The pericentromere features low-complexity tandem repeats (gray) flanking a higher complexity centromeric array (red arrow). Putative centromere locations for both haplotypes of all isolates are provided in Table S16.

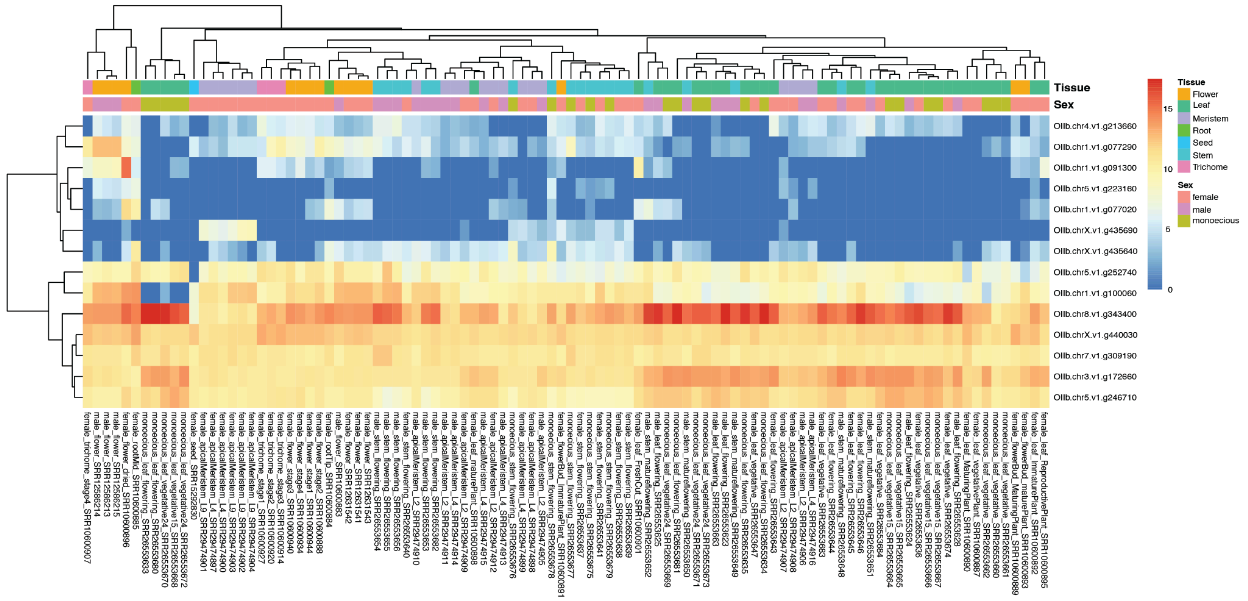


**Figure S10. Heatmap of expression across 100 cannabis RNAseq libraries.** The selected genes here are involved with ethylene, including homologs of aminocyclopropane-1-carboxylate synthase (ACC) synthase (*ACS*: OIIb.chrX.v1.g435690, OIIb.chr4.v1.g213660, OIIb.chr5.v1.g223160, OIIb.chr1.v1.g077020, OIIb.chr1.v1.g091300, OIIb.chr5.v1.g252740, OIIb.chr3.v1.g172660, OIIb.chr5.v1.g246710), ACC oxidase (*ACO*: OIIb.chrX.v1.g435640, OIIb.chr1.v1.g077290, OIIb.chr1.v1.g100060, OIIb.chr8.v1.g343400), and ETHYLENE RESPONSE (ETR: OIIb.chrX.v1.g440030, OIIb.chr7.v1.g309190).

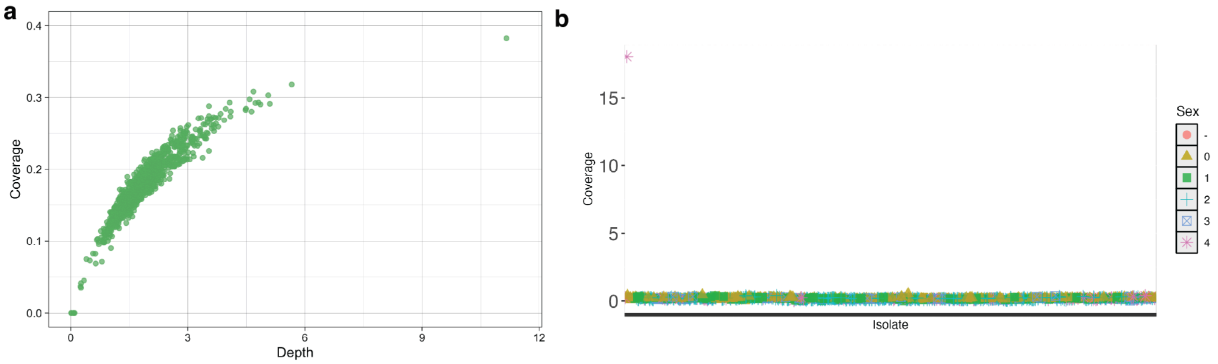


**Fig. S11. Sequencing for the backcross segregating for the monoecy in cannabis. a)** depth and coverage for the 950 samples sequenced for the backcross and controls. **b)** checking for evidence of the Y chromosome in the Khufu data. Samples were sorted by coverage to the MSY (see Materials and Methods for specific details on generating the coverage). No evidence for the Y chromosome was identified in any sample, except for one known male.

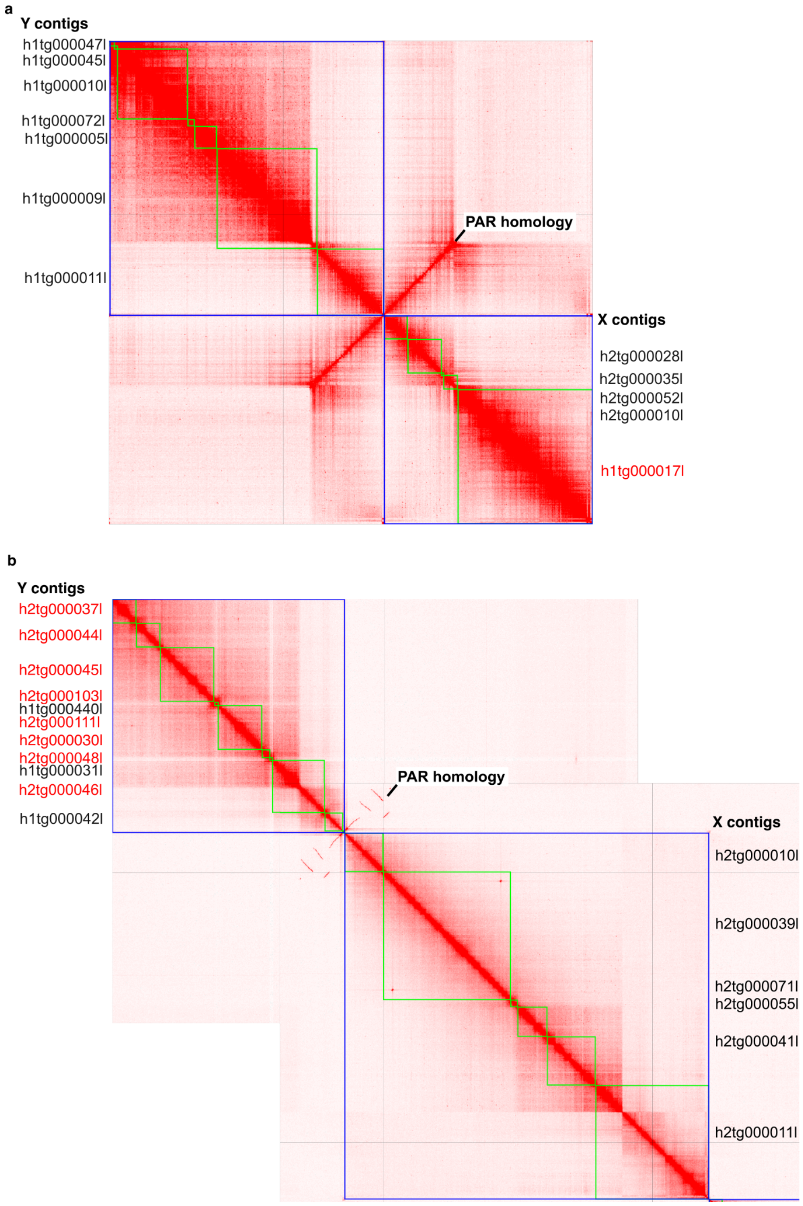


**Fig S12. Hi-C contact map for identifying the sex chromosomes in a joint scaffolded assembly. a**) In the *Cannabis sativa* cv. Carmagnola isolate, the Y-linked contigs identified were all in haplotype 1, but so the contig on the X chromosome that corresponds to the homologous region to the sex-determination region (HXR; red). The HXR contig was manually placed in haplotype 2 with the contigs that correspond to the pseudoautosomal region (PAR). **b**) In the H*umulus lupulus* var. *lupulus* cv. 21110M isolate, The Y chromosome was found to be partially split between both haplotypes; those found in HAP2 are in red and were manually placed in HAP1. This approach was used to validate the Y-linked contigs and identify the X-linked contigs, but was not used for the final scaffolded references.
